# Post-Cyclase Skeletal Rearrangements in Plant Triterpenoid Biosynthesis by a Pair of Branchpoint Isomerases

**DOI:** 10.1101/2022.09.23.508984

**Authors:** Ling Chuang, Shenyu Liu, Jakob Franke

## Abstract

Triterpenoids possess potent biological activities, but their polycyclic skeletons are challenging to synthesise. In biochemistry, the skeletal diversity of plant triterpenoids is normally generated by oxidosqualene cyclases and remains unaltered during subsequent tailoring steps. In contrast, we report here enzyme-mediated skeletal rearrangements after the initial cyclisation, controlling the pathway bifurcation between different plant triterpenoid classes. Using a combination of bioinformatics, heterologous expression in plants and chemical analyses, we identified a cytochrome P450 monooxygenase and two isomerases for this process. The two isomerases share one epoxide substrate but generate two different rearrangement products, one containing a cyclopropane ring. Our findings reveal a new strategy how triterpenoid skeletal diversity is generated in Nature and are crucial for the biotechnological production of limonoid, quassinoid, isoprotolimonoid and glabretane triterpenoids.

## Introduction

Triterpenoids are of great interest to natural product chemists, organic chemists, and medicinal chemists alike due to their complex structures and wide array of bioactivities.^[1, 2]^ The regio- and stereoselective synthesis or modification of such polycyclic and often densely functionalised molecules remains an outstanding challenge and severely hinders drug development of such compounds.^[3–5]^ As an alternative to synthesis, many organisms, particularly plants, possess elaborated biochemical machineries to produce diverse triterpenoids with high selectivity. The biosynthesis of plant triterpenoids is generally divided into two phases: (1) The underlying polycyclic carbon skeleton is generated by an oxidosqualene cyclase; (2) tailoring enzymes introduce specific functionalisations and decorations, e.g. hydroxylations, acylations, and glycosylations, but leave the carbon skeleton unaltered^[6–11]^ An enzymatic way to achieve skeletal modification of already functionalised triterpenoids would be highly desirable to rapidly expand their chemical space. So far, however, no enzyme is known that can achieve such a post-cyclase modification of triterpenoid scaffolds^[9]^ Plants of the order Sapindales are known for their rich diversity of structurally complex triterpenoids. The two best-known groups are limonoids and quassinoids, which include many industrially and ecologically relevant members (Figure 1).^[12–14]^

**Figure 1.**
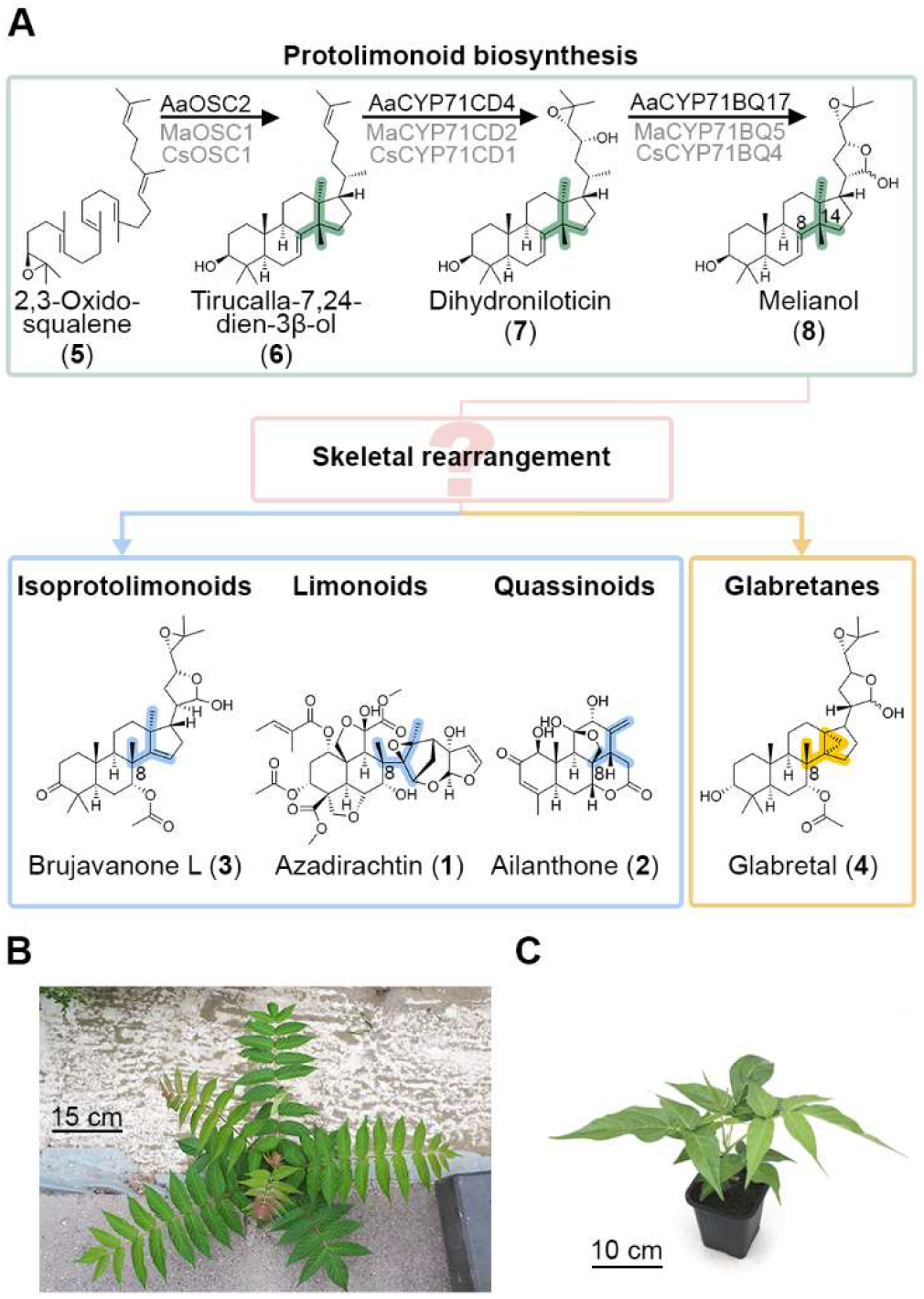
Uncharacterised post-cyclase skeletal rearrangements in plant triterpenoid biosynthesis. A) Plants of the order Sapindales produce different classes of complex triterpenoids, namely isoprotolimonoids, limonoids, quassinoids, and glabretanes, which are derived from the protolimonoid melianol (**8**). Compared to melianol (**8**), downstream triterpenoid classes possess a methyl group at C-8 instead of C-14, and partially also a cyclopropane ring. The enzymes and intermediates involved in these skeletal rearrangements have remained unknown so far. B) and C) The globally invasive plant tree of heaven (Ailanthus altissima, family Simaroubaceae, order Sapindales) was used as a model system in this study.

The limonoid azadirachtin (**1**) is a potent insecticide and key active principle of neem oil.^[14, 15]^ The allelopathic quassinoid ailanthone (**2**) plays a crucial role for the ecological success of the globally invasive tree of heaven (*Ailanthus altissima*), as it occurs in root exudates and helps to outgrow surrounding plants.^[16–18]^ In addition to limonoids and quassinoids, structurally simpler triterpenoids also occur in Sapindales plants, namely compounds related to brujavanone L (**3**) (named isoprotolimonoids herein),^[19]^ and cyclopropane-containing compounds like glabretal (**4**) (named glabretanes herein).^[20]^ Based on increasing structural complexity protolimonoids are considered to be precursors of other triterpenoid classes in Sapindales plants.^[21, 22]^ Protolimonoid biosynthesis requires three steps starting from the common triterpenoid precursor 2,3-oxidosqualene (**5**).^[21, 23]^ First, 2,3-oxidosqualene (**5**) is converted by an oxidosqualene synthase (OSC) into tirucalla-7,24-dien-3β-ol (**6**), followed by multiple oxidations carried out by two cytochrome P450 monooxygenases (CYP450s) which sequentially oxidise **6** to dihydroniloticin (**7**) and then to the protolimonoid melianol (**8**), which is considered to be a key intermediate in these metabolic pathways (Figure 1).^[21, 23]^

Remarkably, the carbon skeletons of protolimonoids exhibit distinct differences to other Sapindales triterpenoid classes, namely the positioning of a methyl group at either C-14 or C-8, and the presence or absence of a cyclopropane ring. In contrast to the general triterpenoid biosynthesis paradigm, this suggests that there are yet unknown enzymes in Sapindales plants which modify the protolimonoid skeleton after the initial cyclisation, leading to pathway bifurcation between the isoprotolimonoids/limonoids/quassinoids groups and the cyclopropane-containing glabretanes. Here we use a combination of co-expression analysis *via* self-organising maps, transient co-expression in the plant host *Nicotiana benthamiana*, and NMR-based structure elucidation to unravel the enzymatic steps and intermediates of this metabolic branchpoint. Three enzymes – a cytochrome P450 monooxygenase and two homologous isomerases evolved from sterol metabolism – are responsible for this post-cyclase skeletal modification and pathway branching en route to biologically active triterpenoids of the limonoid, quassinoid, isoprotolimonoid and glabretane classes and form the basis for future biotechnological approaches.

## Results and Discussion

The accessibility of the tree of heaven (*Ailanthus altissima*) as an invasive plant makes it an ideal model system for studying the biosynthetic pathways of Sapindales triterpenoids. Elucidation of biosynthetic pathways in plants, however, is challenging compared to microbes, as biosynthetic genes are typically not physically clustered. Many recent examples demonstrate that co-expression analysis is a helpful tool to discover novel biosynthetic genes.^[24–27]^ During our recent discovery of the genes required for melianol (**8**) biosynthesis in *A. altissima* based on *de novo* transcriptome sequencing,^[23]^ we observed that these genes were highly co-expressed. We therefore expected that further, yet unknown genes involved in triterpenoid biosynthesis may share a similar expression profile. We therefore searched our *A. altissima* expression data from previous work^[23]^ for gene candidates co-expressed with the first three genes in the pathway. To facilitate visual analysis of the underlying multidimensional dataset, we employed self-organising map (SOM) analysis (Figure 2), which arranges large numbers of transcripts into clusters based on their expression profile.^[28]^ This process successfully grouped the two previously identified cytochrome P450 monooxygenase (CYP450) genes (*AaCYP71CD4* and *AaCYP71BQ17*)^[23]^ into a single cluster with predominant average expression in roots. This cluster was of high quality, i.e. representing a homogeneous group of 695 contigs that was well separated from neighbouring clusters (Figure 2 and Figure S1). The oxidosqualene cyclase gene *AaOSC2* was found in a neighbouring cluster that represented transcripts with high expression in both stem bark and roots. Due to the order of biosynthetic steps, we hypothesised that further pathway genes should have an expression profile more similar to the P450 genes than to the OSC gene and therefore decided to focus on the first cluster for detailed analysis.

**Figure 2.**
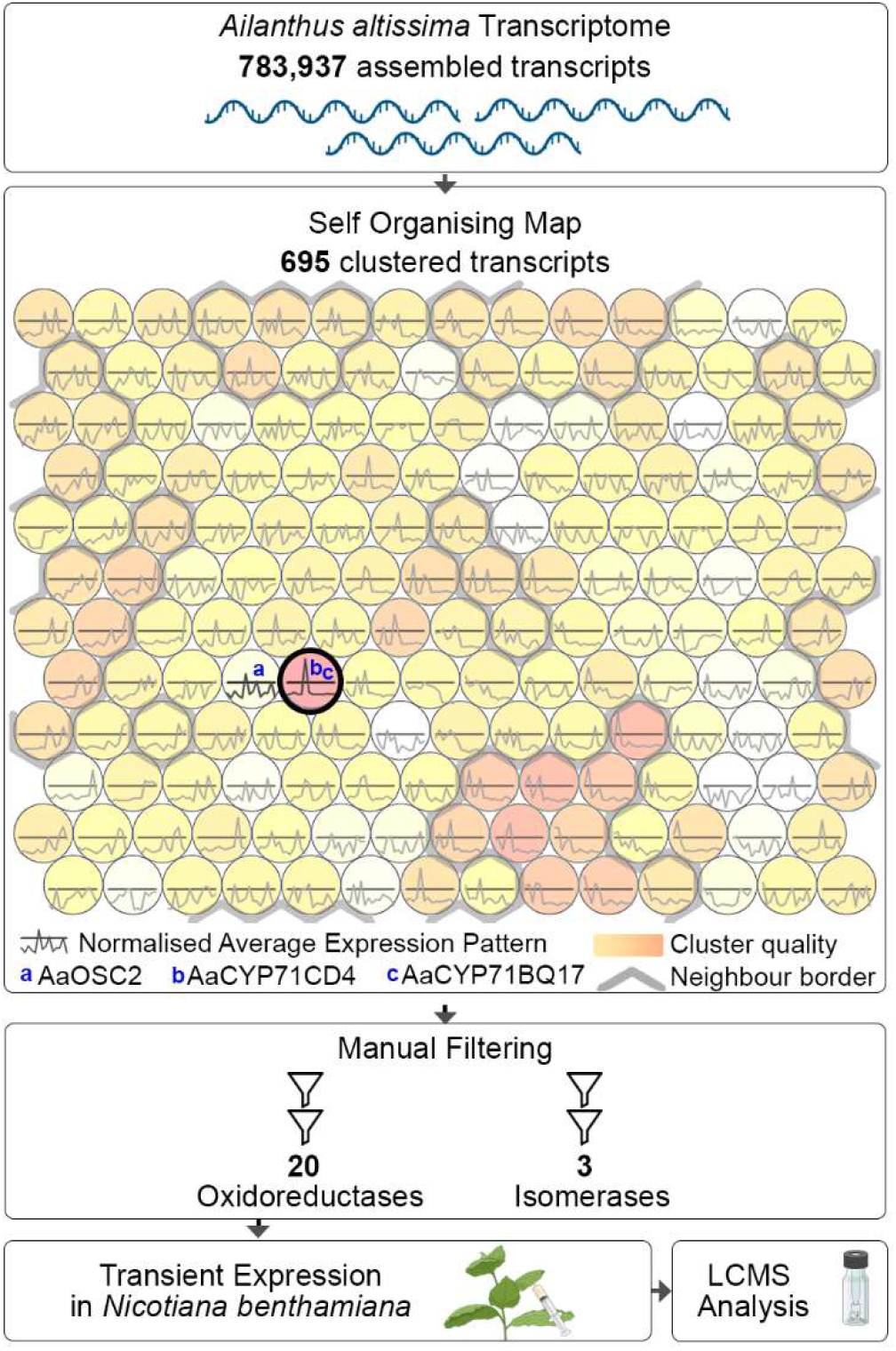
Selection of triterpenoid biosynthesis gene candidates by self-organising map (SOM) analysis. Candidate genes co-expressed with the two pathway genes AaCYP71CD4 and AaCYP71BQ17 were obtained by self-organising map analysis from multidimensional RNA-Seq expression data of Ailanthus altissima transcripts. Both CYP genes were grouped in the same cluster with the highest quality that contained 695 transcripts. The 695 contigs were manually filtered (Figure S2), leading to 20 oxidoreductase and 3 isomerase gene candidates, which were functionally evaluated by transient expression in Nicotiana benthamiana. See main text and supporting information for further details.

It was proposed earlier that the rearrangement of the protolimonoid skeleton could be triggered by epoxidation of the C-7,8 double bond.^[21,22,29]^ To search for the corresponding enzymes, we filtered the 695 contigs obtained by self-organising map analysis for suitable annotations and manually excluded contigs of insufficient length (< 1 kb) and low expression in root (< 50 transcripts per million) (Figure S2). This resulted in a list of 20 oxidoreductase and 3 isomerase gene candidates that were selected for functional screening.

To test if one of these 20 oxidoreductase candidates uses melianol (**8**) as a substrate, we co-expressed the gene candidates with the other pathway genes *AaOSC2, AaCYP71CD4* and *AaCYP71BQ17* as well as booster genes to increase levels of the precursor 2,3-oxidosqualene (**5**) in the plant host *Nicotiana benthamiana. N. benthamiana* is a popular tool for elucidating plant biosynthetic pathways thanks to the capacity to rapidly co-express multiple gene candidates without the need for multiple selection markers[^27,30,31]^ 18 of the 20 oxidase gene candidates were successfully cloned into *Agrobacterium tumefaciens* and co-infiltrated with *AaOSC2, AaCYP71CD1* and *AaCYP71BQ1* into *N. benthamiana*. Crude extracts of the co-expressing plants were then analysed by LC-MS to look for consumption of melianol (**8**) and the production of new compounds. Gratifyingly, co-expression of one candidate gene *(AaCYP88A154*), encoding a cytochrome P450 monooxygenase, showed a clear decrease of melianol (**8**) compared to controls and other samples (Figure 3), whereas all other candidates showed no or inconsistent activity on melianol (**8**) or its precursors **6** or **7**. Two major new peaks (compound **9** at 8.0 min, compound **10** at 6.8 min), and a minor peak at 5.5 min were observed. The new peaks showed putative molecular ions of *m/z* 511 ([M+Na]^+^). In comparison to melianol (m/z 495 for [M+Na]^+^), this implied incorporation of an additional oxygen atom. We suspected that one of the new products might be the previously postulated epoxide of melianol (**8**), while the others could be rearrangement, degradation, or shunt products. To support this hypothesis, we treated melianol (**8**) with *meta*-chloroperoxybenzoic acid (*m*-CPBA), a common epoxidation reagent.^[32]^ Indeed, analysis of the reaction profile showed complete disappearance of **8** and the same three peaks as judged by retention times and mass spectra (Figure 3A). To understand the reaction course, we attempted to purify the two major products from large scale expression in *N. benthamiana*. This was aggravated by the fact that all products were highly sensitive to traces of acid, e.g. from silica, formic acid, or CDCl_3_ (Figures S3, S4, S12). For **9**, switching from CDCl_3_ to C_6_D_6_ as an NMR solvent proved critical to prevent degradation during measurements. We succeeded to purify **9** and **10** as mixtures of lactol epimers and fully elucidate their structures by NMR spectroscopy (Figure 3B). Compound **9** was structurally similar to melianol (**8**), but lacked signals for the C-7,8 double bond. Instead, two new carbon signals at 63.22 and 55.05 for the major epimer suggested the presence of an epoxide at this position. Compound **10** still contained two olefinic carbons, but at C-14,15 instead of C-7,8, and in addition a hydroxy group could be identified at C-7. By detailed 2D NMR analysis, we also identified substructures of two major degradation products present in our NMR sample of **10** formed by opening of the C24/25 epoxide in the side chain (Table S5). Both **9** and **10** are new natural products that we named 7,8-epoxymelianol and isomeliandiol, respectively. Even though isomeliandiol (**10**) has not been found in nature before, ca. 120 natural products with the same carbon skeleton have been isolated from Simaroubaceae, Meliaceae and Rutaceae plants (Table S7), which we name isoprotolimonoids. The presence of an oxygen atom at C-7 and the shift of the methyl group from C-14 to C-8 is also a hallmark feature of mature quassinoids and limonoids.^[22, 33]^ Close homologues of AaCYP88A154 exist in the limonoid-producing plants *Citrus sinensis* (Cs7g22820.1, 85% AA identity) and *Citrus grandis* (CgUng000240.1, 87% AA identity). Hence, in support of earlier biosynthetic proposals,^[22, 29]^ we conclude that the epoxidase AaCYP88A154 catalyses a central step in the biosynthetic pathway of isoprotolimonoids, quassinoids and limonoids, and that 7,8-epoxymelianol (**9**) and isomeliandiol (**10**) are true biosynthetic intermediates that have so far been overlooked due to their instability and rapid conversion.

**Figure 3.**
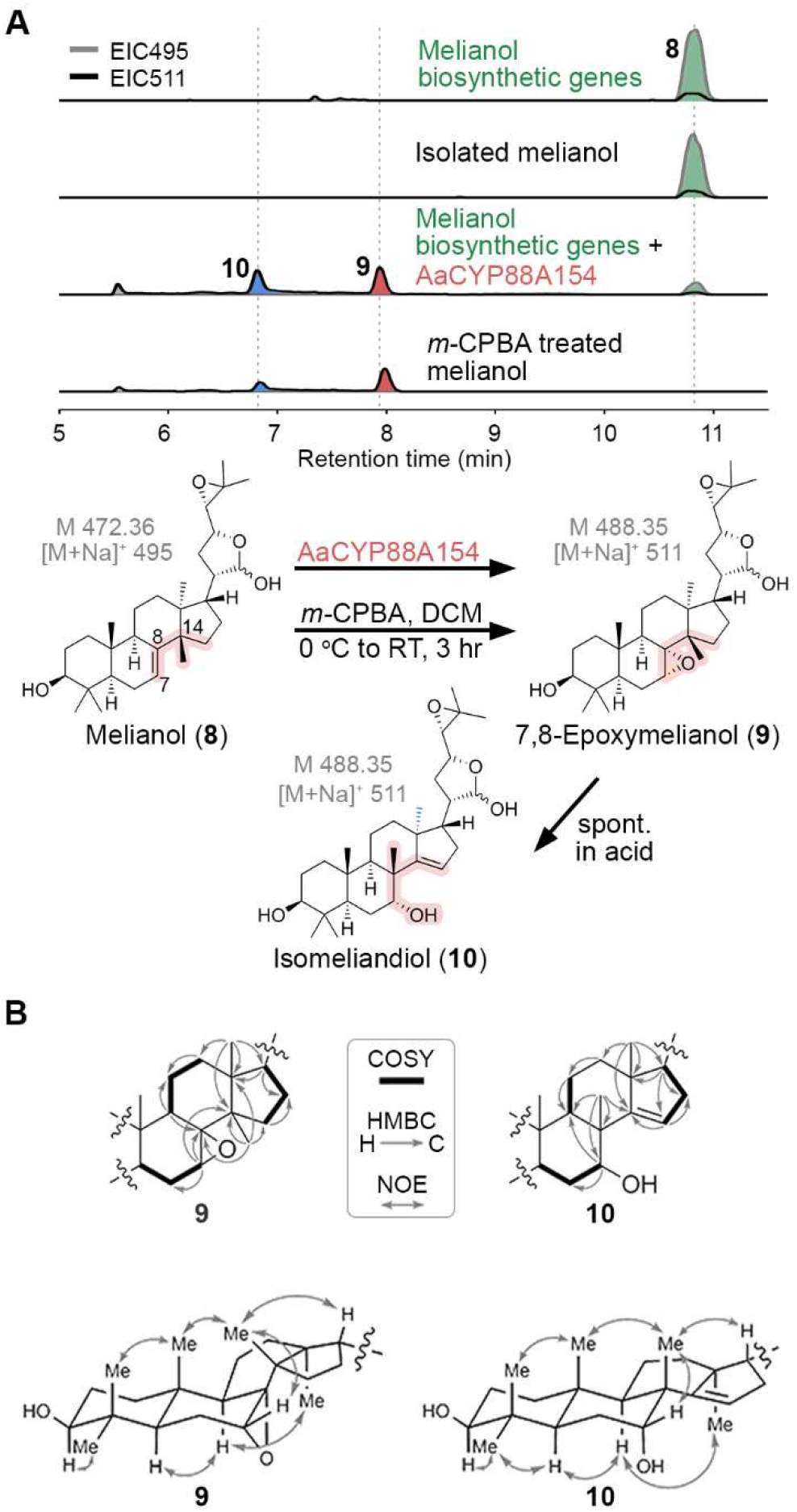
Discovery of the key intermediate 7,8-epoxymelianol (**9**). A) Epoxidation of melianol by AaCYP88A154 or chemically using meta-chloroperoxybenzoic acid (m-CPBA) generates the previously unknown biosynthetic intermediates 7,8-epoxymelianol (**9**) and isomeliandiol (**10**). Melianol was generated in situ by co-expression of melianol biosynthetic genes (AaOSC2, AaCYP71CD4 and AaCYP71BQ17) and booster genes in Nicotiana benthamiana. B) Key COSY, HMBC, and NOE correlations for structure elucidation of **9** and **10**.

Given the high reactivity of 7,8-epoxymelianol (**9**), we speculated that additional enzymes might direct its further conversion *in planta*. The involvement of non-oxidative enzymes such as isomerases for this step has also been proposed previously by Hodgson *et* al.^[21]^ We therefore next focussed on the 3 isomerase candidates from our self-organising map analysis (Figure 2, Figure S2). The isomerase gene candidates were again cloned into a transient expression vector and co-expressed with the other pathway genes *AaOSC2, AaCYP71CD4, AaCYP71BQ17*, and *AaCYP88A154* in *N. benthamiana*. Strikingly, in the presence of candidate ISM1, a strong shift towards isomeliandiol (**10**) as the major product occurred (Figure 4A). To our bigger surprise, presence of candidate ISM2 also led to almost complete disappearance of epoxide **9**, but a new product peak (compound **11**) at 6.6 min was observed. The last candidate did not show any changes to the metabolic profile compared to controls. Like for **9** and **10**, the mass spectrum of **11** showed a molecular ion of *m*/*z* 511, suggesting it to be a previously not observed isomer. We isolated **11** from large scale co-expression in *N. benthamiana*. Strikingly, NMR analysis of **11** clearly showed the presence of a cyclopropane, as judged by a CH2 group with unusual high field shifts (δ_C_ = 13.9 ppm, δ_H_ = 0.65 / 0.45 ppm) (Figure 4B).^[34]^ Full structure elucidation of **11** (Figure 4C) indicated a novel natural product structurally related to glabretal, a triterpenoid previously isolated from *Guarea glabra* (Meliaceae),^[20]^ one of ca. 110 natural products with the same carbon skeleton found in the families Meliaceae, Rutaceae, and Simaroubaceae for which we suggest the name glabretanes (Table S8). Hence, we named **11** protoglabretal. Taken together, our findings show that the two isomerases ISM1 and ISM2 control the skeletal rearrangement cascade of 7,8-epoxymelianol (**9**), leading to two different triterpenoid skeletons. While ISM1 merely channels the spontaneous reaction towards isomeliandiol (**10**), ISM2 generates a product that is not observed when the rearrangement occurs spontaneously. Even though both ISM1 and ISM2 genes have highly similar expression profiles (Figure S1B), co-expression of both genes in *N. benthamiana* demonstrated that isomeliandiol (**10**) and protoglabretal (**11**) can be formed in parallel *in planta* (Figure 4A). We therefore conclude that ISM1 and ISM2 are central gatekeepers in plant triterpenoid metabolism, controlling the formation of the isoprotolimonoid and glabretane structural subclasses.

**Figure 4.**
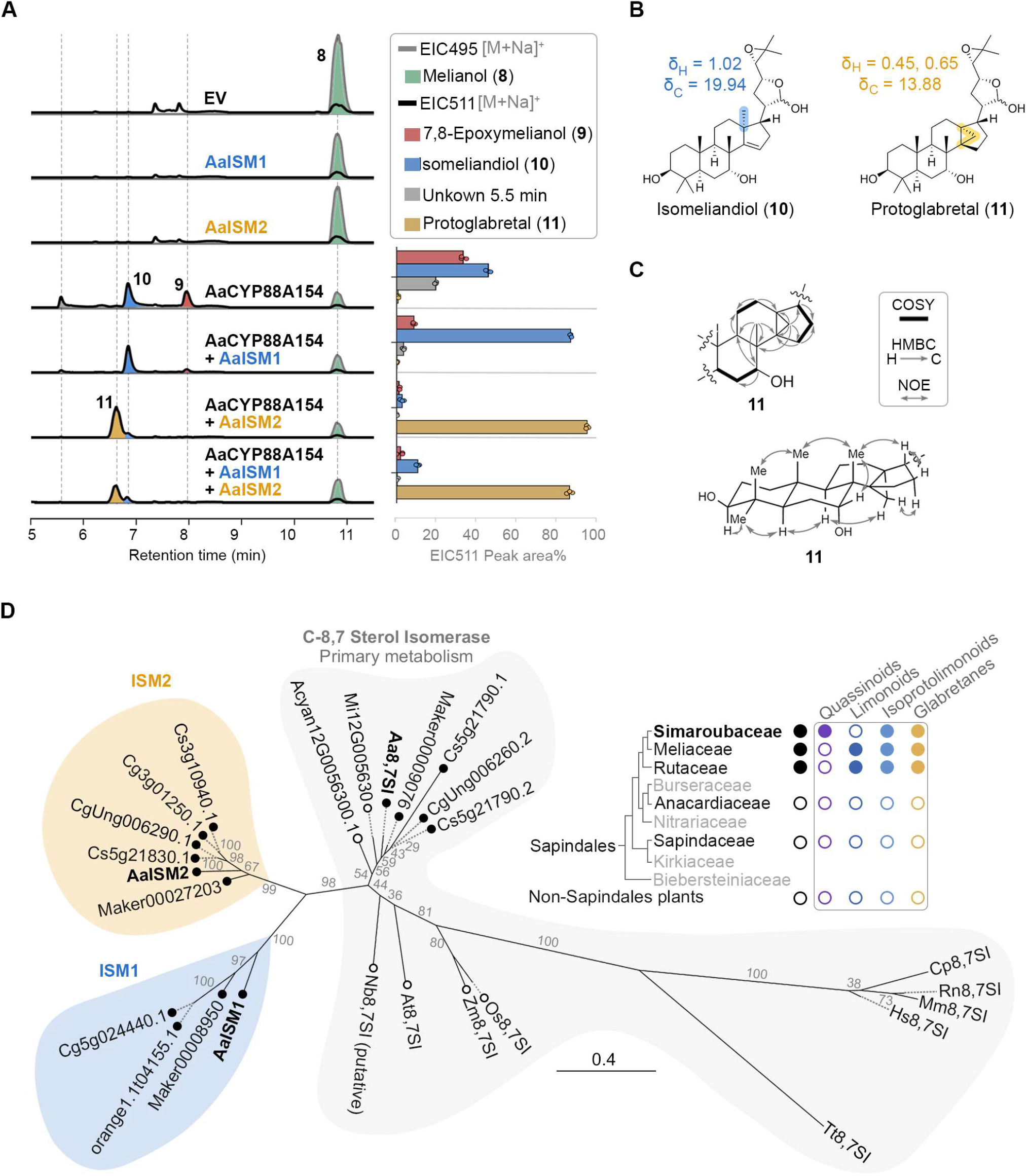
The homologous isomerases AaISM1 and AaISM2 channel the rearrangement of 7,8-epoxymelianol (**9**) towards isomeliandiol (**10**) and protoglabretal (**11**), respectively. (A) LCMS profiles of AaISM1 and AaISM2 genes co-expressed with genes for the biosynthesis of 7,8-epoxymelianol (**9**) in N. benthamiana. The product distribution based on relative peak areas of compounds **9**, **10**, **11** and the minor side product at 5.5 min compared to their total peak area is additionally shown as a bar plot for three biological replicates each. (B) Structures of the new natural products isomeliandiol (**10**) and protoglabretal (**11**) together with chemical shifts at C-18 (highlighted). C) Key COSY, HMBC, and NOE correlations for **11**. (D) Maximum-likelihood phylogenetic tree of sterol isomerases. The scale bar indicates the phylogenetic distance. Bootstrap values for 100 replicates are shown. Homologues of AaISM1 and AaISM2 are found in Meliaceae (Toona sinensis (IDs start with Maker)) and Rutaceae (**C**itrus **s**inensis; **C**itrus **g**randis) plants, but not in plants from other Sapindales families (**Ac**er **yan**gbiense, Sapindaceae; **M**angifera **i**ndica, Anacardiaceae), matching the distribution of quassinoids, limonoids, isoprotolimonoids, and glabretanes in these families. For further details see main text and Supporting Information.

The amino acid identity between ISM1 and ISM2 is 50%. Both are related to C-8,7 sterol isomerases (8,7SI) from primary metabolism, which catalyse the key isomerisation of the Δ^8^ double bond to Δ^7^ in sterol biosynthesis in all eukaryotes (Figure S30).^[35–39]^ The most well-known C-8,7 sterol isomerase from plants is HYDRA1 / At8,7SI from *Arabidopsis thaliana*.^[39, 40]^ The amino acid identity of At8,7SI compared to ISM1 and ISM2 is 45% and 53%, respectively. We generated a phylogenetic tree with other C-8,7 sterol isomerases from primary metabolism as well as putative homologues of ISM1 and ISM2 found in publicly available genome data of the limonoid-producing plants *Citrus sinensis* (sweet orange), *Citrus grandis* (pomelo),^[41, 42]^ and *Toona sinensis*.^[43]^ This analysis suggested that homologues of AaISM1 and AaISM2 are conserved in Rutaceae and Meliaceae but not in other plants, matching the occurrence of quassinoid, limonoid, isoprotolimonoid, and glabretane triterpenoids and thus supporting the proposed key roles of ISM1 and ISM2 for triterpenoid metabolism (Figure 4D). Our phylogenetic analysis suggests that both ISM1 and ISM2 evolved by duplication of a C-8,7 sterol isomerase gene from primary metabolism. Such gene duplication and neofunctionalisation events are known as key drivers for the evolution of plant specialised metabolism.^[44–46]^

Only very few other examples of isomerases in plant specialised metabolism are known, and none performs a skeletal rearrangement or affects more than three adjacent atoms (Figure S31). Most of these examples play a role in non-terpenoid metabolic pathways, namely chalcone isomerase in flavonoid metabolism,^[47, 48]^ the BAHD acyltransferase COSY in coumarin biosynthesis,^[49]^ and neopinone isomerase in opiate production.^[50]^ Two isomerases are known from plant terpenoid metabolism, but only catalyse double bond shifts: In withanolide biosynthesis an isomerase evolved from a reductase performs Δ^24(28)^ to a Δ^24(25)^ double bond isomerisation.^[51]^ A similar double bond shift was reported for Δ^5^-3-ketosteroid isomerase in the context of cardenolide biosynthesis.^[52]^ In contrast to these known isomerases, ISM1 and ISM2 modify the underlying scaffolds of their substrates. Mechanistically, we propose that the preceding epoxidation enables a cationic rearrangement cascade that involves multiple carbon atoms in spatial proximity, typical for terpenoid cyclisations and rearrangements (Figure 5).^[53–55]^ While several ways for enzymatic cyclopropanation are already known,^[56–59]^ ISM2 represents a new way how a cyclopropane can be installed onto an existing triterpenoid skeleton. The fact that the two related enzymes ISM1 and ISM2 generate different rearrangement products from the same epoxide substrate also suggests that protein engineering is highly promising to harness these isomerases for even further skeletal modifications in the future. Homologues of these enzymes from a different model system were also independently discovered by Osbourn, Sattely, and co-workers (manuscript submitted).

**Figure 5.**
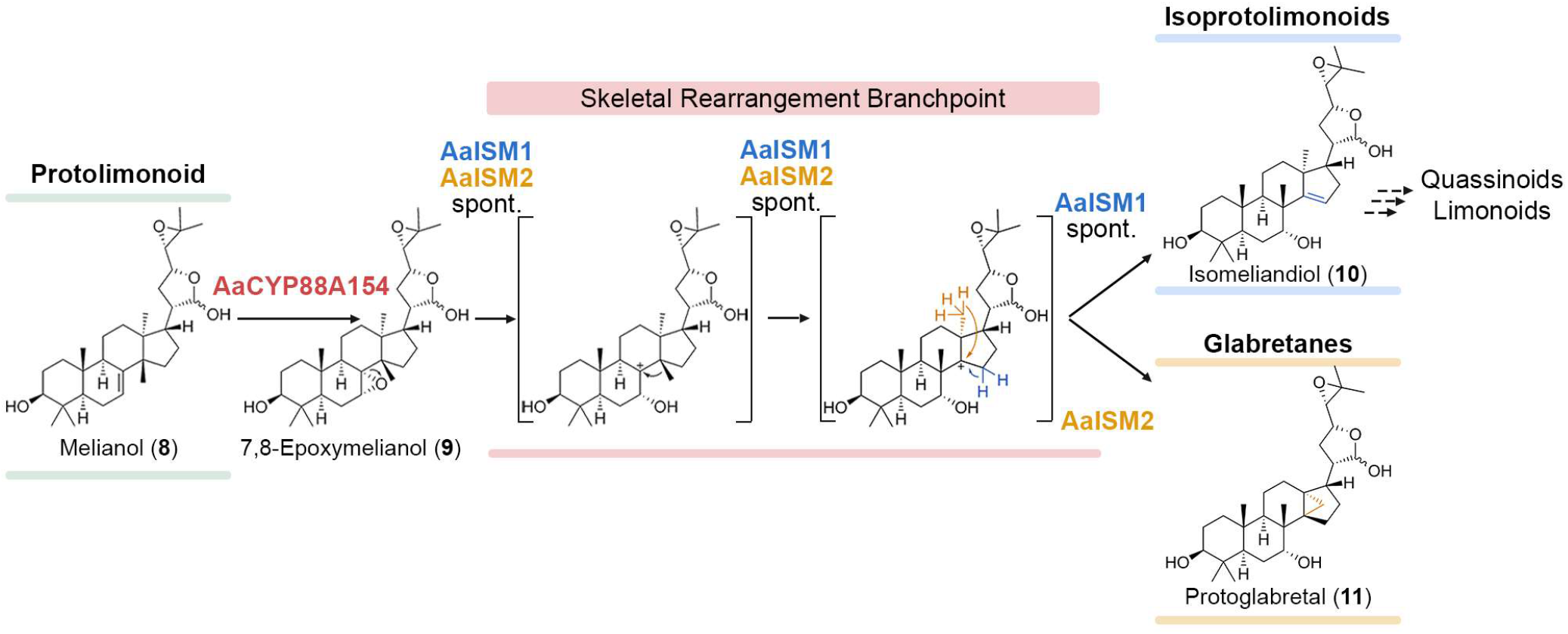
Proposed mechanism for the rearrangement cascade from melianol (**8**). Triggered by C-7,8 epoxidation via the cytochrome P450 monooxygenase AaCYP88A154, the two isomerases AaISM1 and AaISM2 generate the previously unknown triterpenoids isomeliandiol (**10**) and protoglabretal (**11**) by directing the fate of carbon cation intermediates. Thereby, ISM1 and ISM2 control the branching between the isoprotolimonoid/limonoid/quassinoid and glabretane classes of triterpenoids, respectively.

## Conclusion

Taken together, we discovered three novel enzymes in plant triterpenoid metabolism, a cytochrome P450 and two isomerases, that perform post-cyclase skeletal rearrangements of triterpenoids in Sapindales plants at a metabolic branchpoint. The two isomerases AaISM1 and AaISM2 share the same substrate, 7,8-epoxymelianol (**9**), formed by the cytochrome P450 AaCYP88A154, but generate two different rearrangement products isomeliandiol (**10**) and protoglabretal (**11**), representing different classes of triterpenoids. Traditionally, the skeletal diversity of triterpenoids was considered to be solely derived from the initial cyclisation. Our discovery here now shows how triterpenoid skeletal diversity can also be generated beyond the cyclisation phase in Nature. Our findings pave the way for developing new strategies to achieve skeletal editing of already cyclised triterpenoids, which would greatly facilitate the speed by which already functionalised triterpenoids can be generated for medicinal chemistry and other applications. Lastly, the biosynthetic genes described herein will also be crucial to develop biotechnological tools for the production of industrially relevant triterpenoids like the insecticidal limonoid azadirachtin (**1**) and many other triterpenoids belonging to the isoprotolimonoid, limonoid, quassinoid, or glabretane classes.

## Experimental Section

See supporting information.

## Acknowledgements

We thank Prof. Dr. David Nelson (Department of Molecular Science, University of Tennessee, Memphis, USA) and the P450 nomenclature committee for naming AaCYP88A154, Yvonne Leye and Miriam Fent for excellent horticultural support, Katja Körner for excellent technical support, and Johanna Wolf and Yue Sun for assistance with the transient expression screening. We thank Prof. Dr. Christian Hertweck (Leibniz Institute for Natural Product Research and Infection Biology, HKI, Jena, Germany) for helpful discussions. We gratefully acknowledge financial support by the Fonds der Chemischen Industrie, the Emmy Noether programme of the Deutsche Forschungsgemeinschaft (FR 3720/3-1) and the SMART BIOTECS alliance between the Technische Universität Braunschweig and the Leibniz Universität Hannover, supported by the Ministry of Science and Culture (MWK) of Lower Saxony. We also thank the DFG for the provision of NMR equipment (INST 187/686-1). In addition, this work was supported by the LUH compute cluster, which is funded by the Leibniz Universität Hannover, the Lower Saxony Ministry of Science and Culture (MWK) and the German Research Association (DFG).

## References

[1] R. A. Hill, J. D. Connolly, Nat. Prod. Rep. 2020, 37, 962–998.

[2] M. Zhou, R.-H. Zhang, M. Wang, G.-B. Xu, S.-G. Liao, Eur. J. Med. Chem. 2017, 131, 222–236.

[3] S. Yamashita, A. Naruko, Y. Nakazawa, L. Zhao, Y. Hayashi, M. Hirama, Angew. Chem. Int. Ed. 2015, 54, 8538–8541.

[4] L. Furiassi, E. J. Tonogai, P. J. Hergenrother, Angew. Chem. Int. Ed. 2021, 60, 16119–16128.

[5] E. J. Pazur, P. Wipf, Org. Biomol. Chem. 2022, 20, 3870–3889.

[6] R. Thimmappa, K. Geisler, T. Louveau, P. O’Maille, A. Osbourn, Annu. Rev. Plant Biol. 2014, 65, 225–257.

[7] S. Ghosh, Front. Plant Sci. 2017, 8.

[8] S. Sawai, K. Saito, Front. Plant Sci. 2011, 2.

[9] A. Almeida, L. Dong, G. Appendino, S. Bak, Nat. Prod. Rep. 2020, DOI 10.1039/C9NP00030E.

[10] D. Hu, H. Gao, X. Yao, in Compr. Nat. Prod. III (Eds.: H.-W. (Ben) Liu, T.P. Begley), Elsevier, Oxford, 2020, pp. 577–612.

[11] P. D. Cárdenas, A. Almeida, S. Bak, Front. Plant Sci. 2019, 10, 1523.

[12] E. Houël, D. Stien, G. Bourdy, E. Deharo, in Nat. Prod. (Eds.: K.G. Ramawat, J.-M. Mérillon), Springer Berlin Heidelberg, Berlin, Heidelberg, 2013, pp. 3775–3802.

[13] G. Fiaschetti, M. A. Grotzer, T. Shalaby, D. Castelletti, A. Arcaro, Curr. Med. Chem. 2011, 18, 316–328.

[14] Q.-G. Tan, X.-D. Luo, Chem. Rev. 2011, 111, 7437–7522.

[15] S. Fan, C. Zhang, T. Luo, J. Wang, Y. Tang, Z. Chen, L. Yu, Molecules 2019, 24, 3679.

[16] R. M. Heisey, Am. J. Bot. 1996, 83, 192–200.

[17] I. A. B. S. Alves, H. M. Miranda, L. A. L. Soares, K. P. Randau, Rev. Bras. Farmacogn. 2014, 24, 481–501.

[18] B. Sladonja, M. Sušek, J. Guillermic, Environ. Manage. 2015, 56, 1009–1034.

[19] S.-H. Dong, J. Liu, Y.-Z. Ge, L. Dong, C.-H. Xu, J. Ding, J.-M. Yue, Phytochemistry 2013, 85, 175–184.

[20] G. Ferguson, P. A. Gunn, W. C. Marsh, R. McCrindle, R. Restivo, J. D. Connolly, J. W. B. Fulke, M. S. Henderson, J. Chem. Soc. Chem. Commun. 1973, 159–160.

[21] H. Hodgson, R. D. L. Peña, M. J. Stephenson, R. Thimmappa, J. L. Vincent, E. S. Sattely, A. Osbourn, Proc. Natl. Acad. Sci. U. S. A. 2019, 116, 17096–17104.

[22] M. F. Das, G. F. Da Silva, O. R. Gottlieb, Biochem. Syst. Ecol. 1987, 15, 85–103.

[23] L. Chuang, S. Liu, D. Biedermann, J. Franke, Front. Plant Sci. 2022, 13.

[24] L. Caputi, J. Franke, S. C. Farrow, K. Chung, R. M. E. Payne, T.-D. Nguyen, T.-T. T. Dang, I. S. T. Carqueijeiro, K. Koudounas, T. D. de Bernonville, B. Ameyaw, D. M. Jones, I. J. C. Vieira, V. Courdavault, S. E. O’Connor, Science 2018, 360, 1235–1239.

[25] T.-T. T. Dang, J. Franke, I. S. T. Carqueijeiro, C. Langley, V. Courdavault, S. E. O’Connor, Nat. Chem. Biol. 2018, 14, 760–763.

[26] R. S. Nett, Y. Dho, Y.-Y. Low, E. S. Sattely, Proc. Natl. Acad. Sci. U. S. A. 2021, 118, e2102949118.

[27] J. R. Jacobowitz, J.-K. Weng, Annu. Rev. Plant Biol. 2020, 71, null.

[28] R. M. E. Payne, D. Xu, E. Foureau, M. I. S. Teto Carqueijeiro, A. Oudin, T. D. de Bernonville, V. Novak, M. Burow, C.-E. Olsen, D. M. Jones, E. C. Tatsis, A. Pendle, B. A. Halkier, F. Geu-Flores, V. Courdavault, H. H. Nour-Eldin, S. E. O’Connor, Nat. Plants 2017, 3, 16208.

[29] J. Polonsky, in Fortschritte Chem. Org. Naturstoffe Prog. Chem. Org. Nat. Prod. (Eds.: M.J. Cormier, H. Flasch, B. Franck, K. Hori, L. Jaenicke, W. Keller-Schierlein, H.D. Locksley, D.G. Müller, J. Polonsky, R. Tschesche, J.E. Wampler, G. Wulff, H. Grisebach, G.W. Kirby, W. Herz), Springer, Vienna, 1973, pp. 101–150.

[30] L. Chuang, J. Franke, in Eng. Nat. Prod. Biosynth. Methods Protoc. (Ed.: E. Skellam), Springer US, New York, NY, 2022, pp. 395–420.

[31] J. Bally, H. Jung, C. Mortimer, F. Naim, J. G. Philips, R. Hellens, A. Bombarely, M. M. Goodin, P. M. Waterhouse, Annu. Rev. Phytopathol. 2018, 56, 405–426.

[32] H. Hussain, A. Al-Harrasi, I. R. Green, I. Ahmed, G. Abbas, N. Ur Rehman, RSC Adv. 2014, 4, 12882–12917.

[33] D. S. Seigler, in Plant Second. Metab., Springer, Boston, MA, 1998, pp. 473–485.

[34] F. Trottmann, J. Franke, I. Richter, K. Ishida, M. Cyrulies, H.-M. Dahse, L. Regestein, C. Hertweck, Angew. Chem. Int. Ed. 2019, 58, 14129–14133.

[35] T. Long, A. Hassan, B. M. Thompson, J. G. McDonald, J. Wang, X. Li, Nat. Commun. 2019, 10, 2452.

[36] P. Benveniste, Annu. Rev. Plant Biol. 2004, 55, 429–457.

[37] E. Desmond, S. Gribaldo, Genome Biol. Evol. 2009, 1, 364–381.

[38] T. Zhang, D. Yuan, J. Xie, Y. Lei, J. Li, G. Fang, L. Tian, J. Liu, Y. Cui, M. Zhang, Y. Xiao, Y. Xu, J. Zhang, M. Zhu, S. Zhan, S. Li, Mol. Biol. Evol. 2019, 36, 2548–2556.

[39] R. J. Grebenok, T. E. Ohnmeiss, A. Yamamoto, E. D. Huntley, D. W. Galbraith, D. Della Penna, Plant Mol. Biol. 1998, 38, 807–815.

[40] M. Souter, J. Topping, M. Pullen, J. Friml, K. Palme, R. Hackett, D. Grierson, K. Lindsey, Plant Cell 2002, 14, 1017–1031.

[41] Q. Xu, L.-L. Chen, X. Ruan, D. Chen, A. Zhu, C. Chen, D. Bertrand, W.-B. Jiao, B.-H. Hao, M. P. Lyon, J. Chen, S. Gao, F. Xing, H. Lan, J.-W. Chang, X. Ge, Y. Lei, Q. Hu, Y. Miao, L. Wang, S. Xiao, M. K. Biswas, W. Zeng, F. Guo, H. Cao, X. Yang, X.-W. Xu, Y.-J. Cheng, J. Xu, J.-H. Liu, O. J. Luo, Z. Tang, W.-W. Guo, H. Kuang, H.-Y. Zhang, M. L. Roose, N. Nagarajan, X.-X. Deng, Y. Ruan, Nat. Genet. 2012, 45, 59–66.

[42] G. A. Wu, S. Prochnik, J. Jenkins, J. Salse, U. Hellsten, F. Murat, X. Perrier, M. Ruiz, S. Scalabrin, J. Terol, M. A. Takita, K. Labadie, J. Poulain, A. Couloux, K. Jabbari, F. Cattonaro, C. Del Fabbro, S. Pinosio, A. Zuccolo, J. Chapman, J. Grimwood, F. R. Tadeo, L. H. Estornell, J. V. Muñoz-Sanz, V. Ibanez, A. Herrero-Ortega, P. Aleza, J. Pérez-Pérez, D. Ramón, D. Brunel, F. Luro, C. Chen, W. G. Farmerie, B. Desany, C. Kodira, M. Mohiuddin, T. Harkins, K. Fredrikson, P. Burns, A. Lomsadze, M. Borodovsky, G. Reforgiato, J. Freitas-Astúa, F. Quetier, L. Navarro, M. Roose, P. Wincker, J. Schmutz, M. Morgante, M. A. Machado, M. Talon, O. Jaillon, P. Ollitrault, F. Gmitter, D. Rokhsar, Nat. Biotechnol. 2014, 32, 656–662.

[43] Y.-T. Ji, Z. Xiu, C.-H. Chen, Y. Wang, J.-X. Yang, J.-J. Sui, S.-J. Jiang, P. Wang, S.-Y. Yue, Q.-Q. Zhang, J. Jin, G.-S. Wang, Q.-Q. Wei, B. Wei, J. Wang, H.-L. Zhang, Q.-Y. Zhang, J. Liu, C.-J. Liu, J.-B. Jian, C.-Q. Qu, Mol. Ecol. Resour. 2021, 21, 1243–1255.

[44] P. D. Sonawane, A. Jozwiak, S. Panda, A. Aharoni, Curr. Opin. Plant Biol. 2020, 55, 118–128.

[45] A. Jozwiak, P. D. Sonawane, S. Panda, C. Garagounis, K. K. Papadopoulou, B. Abebie, H. Massalha, E. Almekias-Siegl, T. Scherf, A. Aharoni, Nat. Chem. Biol. 2020, 1–9.

[46] T. Tohge, A. R. Fernie, Plants 2020, 9, 622.

[47] Y. Yin, X. Zhang, Z. Gao, T. Hu, Y. Liu, Mol. Biotechnol. 2019, 61, 32–52.

[48] J. M. Jez, M. E. Bowman, R. A. Dixon, J. P. Noel, Nat. Struct. Biol. 2000, 7, 786–791.

[49] R. Vanholme, L. Sundin, K. C. Seetso, H. Kim, X. Liu, J. Li, B. D. Meester, L. Hoengenaert, G. Goeminne, K. Morreel, J. Haustraete, H.-H. Tsai, W. Schmidt, B. Vanholme, J. Ralph, W. Boerjan, Nat. Plants 2019, 5, 1066–1075.

[50] M. Dastmalchi, X. Chen, J. M. Hagel, L. Chang, R. Chen, S. Ramasamy, S. Yeaman, P. J. Facchini, Nat. Chem. Biol. 2019, 15, 384–390.

[51] E. Knoch, S. Sugawara, T. Mori, C. Poulsen, A. Fukushima, J. Harholt, Y. Fujimoto, N. Umemoto, K. Saito, Proc. Natl. Acad. Sci. U. S. A. 2018, 115, E8096–E8103.

[52] N. Meitinger, D. Geiger, T. W. Augusto, R. Maia de Pádua, W. Kreis, Phytochemistry 2015, 109, 6–13.

[53] D. W. Christianson, Chem. Rev. 2017, 117, 11570–11648.

[54] J. S. Dickschat, Angew. Chem. Int. Ed. 2019, 58, 15964–15976.

[55] J. S. Dickschat, Nat. Prod. Rep. 2016, 33, 87–110.

[56] S. Ma, D. Mandalapu, S. Wang, Q. Zhang, Nat. Prod. Rep. 2022, 39, 926–945.

[57] C. J. Thibodeaux, W. Chang, H. Liu, Chem. Rev. 2012, 112, 1681–1709.

[58] L. A. Wessjohann, W. Brandt, T. Thiemann, Chem. Rev. 2003, 103, 1625–1648.

[59] F. Trottmann, K. Ishida, M. Ishida-Ito, H. Kries, M. Groll, C. Hertweck, Nat. Chem. 2022, 14, 884–890.

