## Supplementary Information for "Post-Cyclase Skeletal Rearrangements in Plant Triterpenoid Biosynthesis by a Pair of Branchpoint Isomerases"

Ling Chuang[+], Shenyu Liu[+] and Prof. Dr. Jakob Franke*

### Table of Contents

Table of Contents ... 2
Experimental Procedures ... 3
General chemical methods ... 3
General plant methods ... 3
Self-organising map analysis and candidate selection ... 3
Construction of plasmids for transient expression in *Nicotiana benthamiana* ... 3
Transient expression of candidate genes in *Nicotiana benthamiana* ... 4
Metabolite extraction and LC-MS analysis ... 4
Synthesis of 7,8-epoxymelianol (9) ... 4
Purification of isomeliandiol (10) and protoglabretal (11) ... 5
Isomeliandiol (10): ... 5
Phylogenetic analysis ... 5
References ... 42

**Figures**

**Figure S1. Comparison of average expression patterns of the two clusters selected from SOM analysis to pathway gene expression patterns.** ... 6
**Figure S2. Candidate selection from self-organising map (SOM) analysis of *Ailanthus altissima* transcriptome data.** ... 6
**Figure S3. Acid sensitivity of 7,8-epoxymelianol (9) to silica.** ... 8
**Figure S4. Time course showing degradation of 7,8-epoxymelianol (9) (peak at 6.6 min) in 50/50 MeCN/$H_2O$ + 0.1% formic acid.** ... 8
**Figure S5. $^1$H spectrum of 7,8-epoxymelianol (9) ($C_6D_6$, 298 K, 600 MHz).** ... 14
**Figure S6. $^{13}$C spectrum of 7,8-epoxymelianol (9) ($C_6D_6$, 298 K, 151 MHz).** ... 15
**Figure S7. HSQC spectrum of 7,8-epoxymelianol (9) ($C_6D_6$, 298 K, 600 MHz).** ... 16
**Figure S8. HMBC spectrum of 7,8-epoxymelianol (9) ($C_6D_6$, 298 K, 600 MHz).** ... 17
**Figure S9. COSY spectrum of 7,8-epoxymelianol (9) ($C_6D_6$, 298 K, 600 MHz).** ... 18
**Figure S10. NOESY spectrum of 7,8-epoxymelianol (9) ($C_6D_6$, 298 K, 500 MHz).** ... 19
**Figure S11. $^1$H NMR spectrum of isomeliandiol (10) ($CDCl_3$, 298 K, 500 MHz).** ... 20
**Figure S12. $^{13}$C NMR spectrum of isomeliandiol (10) containing the C24/25 epoxide degradation products 12 and 13 ($CDCl_3$, 298 K, 126 MHz).** ... 21
**Figure S13. HSQC spectrum of isomeliandiol (10) ($CDCl_3$, 298 K, 500 MHz).** ... 22
**Figure S14. HMBC spectrum of isomeliandiol (10) ($CDCl_3$, 298 K, 500 MHz).** ... 23
**Figure S15. COSY spectrum of isomeliandiol (10) ($CDCl_3$, 298 K, 500 MHz).** ... 24
**Figure S16. NOESY spectrum of isomeliandiol (10) ($CDCl_3$, 298 K, 500 MHz).** ... 25
**Figure S17. $^1$H NMR spectrum of protoglabretal (11) ($CDCl_3$, 298 K, 500 MHz).** ... 26
**Figure S18. $^{13}$C NMR spectrum of protoglabretal (11) ($CDCl_3$, 298 K, 126 MHz).** ... 27
**Figure S19. DEPT-135 spectrum of protoglabretal (11) ($CDCl_3$, 298 K, 100 MHz).** ... 28
**Figure S20. HSQC spectrum of protoglabretal (11) ($CDCl_3$, 298 K, 500 MHz).** ... 29
**Figure S21. HMBC spectrum of protoglabretal (11) ($CDCl_3$, 298 K, 500 MHz).** ... 30
**Figure S22. COSY spectrum of protoglabretal (11) ($CDCl_3$, 298 K, 500 MHz).** ... 31
**Figure S23. NOESY spectrum of protoglabretal (11) ($CDCl_3$, 298 K, 500 MHz).** ... 32
**Figure S24. Selected key HMBC and COSY correlations of 7,8-epoxymelianol (9).** ... 33
**Figure S25. Selected NOE correlations of 7,8-epoxymelianol (9).** ... 33
**Figure S26. Selected key HMBC and COSY correlations of isomeliandiol (10).** ... 33
**Figure S27. Selected NOE correlations of isomeliandiol (10).** ... 34
**Figure S28. Selected key HMBC and COSY correlations of protoglabretal (11).** ... 34
**Figure S29. Selected NOE correlations of protoglabretal (11).** ... 34
**Figure S30. Overview over common C-8,7 sterol isomerase (8,7SI) reactions in primary metabolism.** ... 40
**Figure S31. Overview over other isomerase-catalysed reactions in plant specialised metabolism.**[59–63] ... 41

**Tables**

**Table S1. Sequences of the primers used in this study.** ... 7
**Table S2. Conditions for extraction and flash chromatography purification of isomeliandiol (10) and protoglabretal (11).** ... 9
**Table S3. $^1$H and $^{13}$C NMR data of 7,8-epoxymelianol (9) ($C_6D_6$, 298 K, 600 MHz).** ... 10
**Table S4. $^1$H and $^{13}$C NMR data of isomeliandiol (10) obtained as a mixture of epimers ($CDCl_3$, 298 K, 500 MHz).** ... 11
**Table S5. Partial $^1$H and $^{13}$C NMR data of degradation products 12 and 13 present in NMR samples of isomeliandiol (10) ($CDCl_3$, 298 K, 500 MHz).** ... 12
**Table S6. $^1$H and $^{13}$C NMR data of protoglabretal (11) ($CDCl_3$, 298 K, 500 MHz).** ... 13
**Table S7. Representative examples of isoprotolimonoids previously isolated from Meliaceae, Rutaceae, and Simaroubaceae plants out of ca. 120 structurally related natural products listed in Reaxys.** ... 35
**Table S8. Representative examples of glabretanes previously isolated from Meliaceae, Rutaceae, and Simaroubaceae plants out of ca. 110 structurally related natural products listed in Reaxys.** ... 37

#### Experimental Procedures

##### General chemical methods

NMR spectra were recorded using Bruker Ultrashield 400, Ultrashield 500 or Ascend 600 MHz spectrometers operating at 400, 500 and 600 MHz for $^1$H NMR and at 100, 126 and 151 MHz for $^{13}$C NMR. $CDCl_3$ and $C_6D_6$ were used as solvents. Chemical shifts were referenced relative to the residual solvent signals ($CDCl_3$: $\delta_H$ = 7.26 ppm, $\delta_C$ = 77.16 ppm; $C_6D_6$: 7.16 ppm, $\delta_C$ = 128.06 ppm) and expressed in δ values (ppm), with coupling constants reported in Hz. Analysis was conducted with TopSpin (Version 4.0.6) or MestReNova (Version 14.2).
HRMS measurements were carried out on a Waters Alliance 2695 HPLC coupled to a Micromass LCT Premier mass spectrometer.
Analytical and semipreparative LCMS analyses were performed on an Agilent Infinity II 1260 system consisting of a G7167A autosampler, G7116A column thermostat, G7111B quaternary pump, G7110B make-up pump, G7115A diode array detector, G1364F fraction collector, and G6125B single quadrupole mass spectrometer equipped with an ESI source (positive mode, 4000 V, 12 L/min drying gas, 350 °C gas temperature). The columns and gradients used are described below.

Automated flash chromatography was performed on a Biotage Isolera One with the stationary phases, solvents and gradients described below.
Melianol as a reference compound and substrate was obtained by transient expression in *N. benthamiana* followed by purification as described previously.[1] All other reagents were purchased from Sigma-Aldrich, Fisher Scientific and Carl Roth. All reagents were directly used without further purification unless mentioned otherwise. Dry solvents were obtained from Acros Organics and stored under $N_2$ with molecule sieve (3 Å).

##### General plant methods

*Ailanthus altissima* plants were grown from seedlings as described previously.[1] Work with *Ailanthus altissima* in our research group is granted by permit DE-NI-2019-001 (NLWKN, Niedersächsischer Landesbetrieb für Wasserwirtschaft, Küsten- und Naturschutz) in addition to regulation (EU) No. 1143/2014. *Nicotiana benthamiana* LAB strain[2] was grown from seeds in a greenhouse with 11 to 16 hours illumination per day and at a temperature between 21 °C to 23 °C as described previously.[3]

##### Self-organising map analysis and candidate selection

Self-organising map (SOM) analysis was based on expression data from our previously published de novo transcriptome data, covering 14 different tissue samples from *Ailanthus altissima* seedlings or 3-year-old trees.[1] Shortly, raw reads from RNA-Seq were assembled *de novo* with Trinity 2.11.0,[4] and contig expression was determined by Salmon 1.3.0[5] and normalised using the TMM method.[6] The SOM analysis was conducted in R version 4.1.2 using the kohonen package,[7] using a slightly adjusted version of the script reported by Payne *et al*.[8] Cluster quality was calculated based on the within-node distance and inter-nodal distance as described in Payne *et al.*;[8] high cluster quality corresponds to low within-node and low inter-nodal distance. Broad grey lines additionally indicate neighbouring clusters with low inter-nodal distance (25% quantile of all).
The node containing transcripts for the previously reported quassinoid pathway genes *AaCYP71CD4* and *AaCYP71BQ17* was manually screened to select gene candidates. For the 695 contigs in the node, protein sequences were deduced using TransDecoder 5.3.0[9] and protein domains annotated using a search with hmmscan 3.2[10] against the Pfam-A database.[11] Candidates were then extracted by filtering the 695 co-expressed contigs for Pfam, blastx and blastn hits containing the keywords p450, oxidoreductase, oxidase, oxygenase, reductase, dehydrogenase (138 hits). The further filtering steps are shown in Figure S2. An analogous strategy was used for isomerase selection (keywords: isomerase, epimerase, mutase) as shown in Figure S2.

##### Construction of plasmids for transient expression in *Nicotiana benthamiana*

Primers for the amplification of full coding sequences were designed based on transcript sequences of candidate genes. The primer sequences are listed in Table S1. The insert sequences were amplified with SuperFi II polymerase (Thermo Fisher Scientific) from cDNA of seedling root. Amplicons were cloned either into the vector pEAQ-HT[12] by In-Fusion HD cloning (Takara Bio), or into the vector pHREAC[13] as described previously by Golden Gate cloning.[3] Cloned sequences were confirmed by Sanger sequencing. Coding sequences of genes *AaCYP88A154*, *AaISM1*, and *AaISM2* identified in this study were deposited in GenBank under the accession numbers ON942227-ON942229.

##### Transient expression of candidate genes in *Nicotiana benthamiana*

For agroinfiltration into *N. benthamiana*, a procedure described earlier by us was used.[3] In short, the plasmids containing gene candidates for transient expression were transformed into *Agrobacterium tumefaciens* GV3101 by electroporation. *A. tumefaciens* strains were precultured for 2 days at 28 °C in LB + 25 µg/mL gentamicin + 50 µg/mL rifampicin + 50 µg/mL kanamycin. After the preculture, *A. tumefaciens* cells were harvested and resuspended in MMA infiltration buffer (10 mM $MgCl_2$, 10 mM 2-(*N*-morpholino)ethanesulfonic acid (MES), 100 µM acetosyringone). Strains carrying candidate genes were mixed with *A. tumefaciens* strains carrying *AstHMGR* (KY284573) [14], *AaOSC2* (ON595696), *AaCYP71CD4* (ON595698), and *AaCYP71BQ17* (ON595699) (all strains at $OD_{600}$ 0.1) prior to syringe infiltration into the abaxial side of *Nicotiana benthamiana* leaves. After infiltration, plants were maintained in a greenhouse until further analysis. For screening, at least three biological replicates were used for each combination.

##### Metabolite extraction and LC-MS analysis

Around 10 mg dry weight of infiltrated leaves were used for metabolite extraction. Infiltrated leaves were harvested 7 days after infiltration. Five leaf disks were harvested using cork-borer no. 5 (10mm) and lyophilized before extraction with 800 µL 90% methanol. After removal of solid debris by centrifugation, the crude extract was directly used for LC-MS analysis. Samples were separated on a C18 column (Poroshell 120 EC-C18, dimensions: 100 × 4.6 mm, particle size: 2.7 µm) using an LCMS system from Agilent (Agilent 1260 II Infinity, Santa Clara, CA, USA). The column temperature was set at 50 °C. As mobile phase, solvent A (water with 0.1%(v/v) formic acid) and solvent B (acetonitrile with 0.1%(v/v) formic acid) were used. Separation was achieved using the following gradient at a flow rate of 1 mL/min: 0-1 min, 10-40% B; 1-11 min, 40-90% B; 11-13 min, 90% B; 13-13.1 min, 90-10% B; 13.1-15 min, 10% B.

##### Synthesis of 7,8-epoxymelianol (9)

*m*-CPBA was purified before use following Kazmaier's report.[15] The exact concentration was determined by NMR. Melianol (11 mg, 23 µmol, 1 eq.) was charged in a flame-dried Schlenk round bottom flask under $N_2$ atmosphere and dissolved in dry DCM (6 mL). The reaction mixture was cooled to 0 °C with an ice-water bath, and *m*-CPBA (14 mg, 85% purity, 70 µmol, 3 eq.) in dry DCM (4 mL) was added dropwise over 10 mins at 0 °C. When the addition of *m*-CPBA was complete, the reaction mixture was allowed to warm to room temperature and stirred for additional 3 hrs. After that, 5 mL $Na_2SO_3$ solution (5% w/v) was added into the reaction mixture. The biphasic mixture was stirred vigorously for 5 mins. The organic layer was then collected, washed with sat. $NaHCO_3$ (5 mL), water (5 mL) and brine (5 mL) sequentially and dried over $Na_2SO_4$. The solvent was removed with a gentle stream of of $N_2$ to give 7,8-epoxymelianol (**9**) (10 mg, 90%) as white powder. No further purification was performed. For the success of this reaction, a low concentration of substrate proved critical to minimise overoxidation and rearrangements.

**7,8-Epoxymelianol (9):**
HR-ESI-MS: $[M+Na]^+$ = 511.3409 (calcd. For $C_{30}H_{48}O_4Na^+$ 511.3399). $^1H$ and $^{13}C$ NMR data see Table S3.

##### Purification of isomeliandiol (10) and protoglabretal (11)

For compound isolation, *N. benthamiana* plants were vacuum infiltrated in a 9.2 L ROTILABO desiccator (Carl Roth, Karlsruhe, Germany) connected to a MZ 2 NT membrane pump (Vacuubrand, Wertheim, Germany) at 30 mbar for 1.5 min. 60 plants were each used for purification of the AaCYP88A154 and AaISM2 products.
Leaves were harvested 7 days post infiltration and lyophilized for 2-3 days until the dry weight remained constant. The crude plant material was ground at room temperature in a blender to powder form and extracted with ethyl acetate (AaCYP88A154 product) or 90/10 MeOH/$H_2O$ (AaISM2 product), filtered and concentrated *in vacuo*. The crude extracts were purified by successive rounds of flash chromatography (Biotage Isolera One) as described in Table S2. This process yielded 2 mg isomeliandiol (**10**) (0.12 mg / g dry weight) as a white powder, and 22 mg protoglabretal (**11**) (1.3 mg / g dry weight) as a yellow solid.

###### Isomeliandiol (10):

HR-ESI-MS: $[M+Na]^+$ = 511.3389 (calcd. For $C_{30}H_{48}O_4Na^+$ 511.3399). $^1H$ and $^{13}C$ NMR data see Table S4.

###### Protoglabretal (11):

HR-ESI-MS: $[M+Na]^+$ = 511.3380 (calcd. For $C_{30}H_{48}O_4Na^+$ 511.3399). $^1H$ and $^{13}C$ NMR data see Table S6.

##### Phylogenetic analysis

To identify homologues of C-8,7 sterol isomerase (8,7SI), AaISM1 and AaISM2 in Sapindales plants, the peptide sequences of characterised *Arabidopsis thaliana* 8,7SI (AF030357.1)[16], AaISM1 and AaISM2 were used as reference for blastp searches against publicly available databases. For orange (*Citrus sinensis*) and pomelo (*Citrus grandis*) from Rutaceae family, the peptide sequences were downloaded from Citrus Pan-genome to Breeding Database (CPBD) v2.0 and v1.0 respectively[17,18]. The sequences of heiyouchun (*Toona sinensis*) from Meliaceae were downloaded from China National GeneBank DataBase (CNGBdb) under the project ID CNP0000958[19]. Sequences of mango (*Mangifera indica*) from Anacardiaceae family were kindly provided by Wang *et al.*,[20] and yangbi mapel (*Acer yangbiense*) data in Sapindaceae family were downloaded from GigaDB [21]. For plants outside of Sapindales, the model plants *Arabidopsis thaliana* and *Nicotiana benthamiana* were chosen and their sequences were downloaded from TAIR (Araport 11) and Solgenomics (genome sequence v1.0.1) respectively. Unique full length peptide sequences with the best E-values and coverage resulting from the blastp search were chosen for alignment and tree construction.
In addition to At8,7SI, characterized 8,7SI from other plants and non-plants were included as reference. For plants, Zm8,7SI (AY533175.1) from corn (*Zea mays*) [22] and a putative 8,7SI (AK059848.1) from *Oryza sativa* validated at transcript level were included[22,23]. For organisms from other kingdoms, Rn8,7SI(AF071501.1) from rat (*Rattus norvegicus*) [24], Mm8,7SI (X97755.1) from house mouse (*Mus musculus*) [25], Cp8,7SI (Q60490) from Guinea pig (*Cavia porcellus*), Hs8,7SI (NP_006570) from human (*Homo sapiens*) [26,27], and a fungal 8,7SI from *Thermothelomyces thermophilus* (XP_003665660)[28] were included.
A multiple sequence alignment of all isomerases was generated using Clustal Omega 1.2.2[29–31]. Then, a phylogenetic tree was constructed using the maximum likelihood method with PhyML 3.3.20180621[32] in Geneious 2021.1.1.

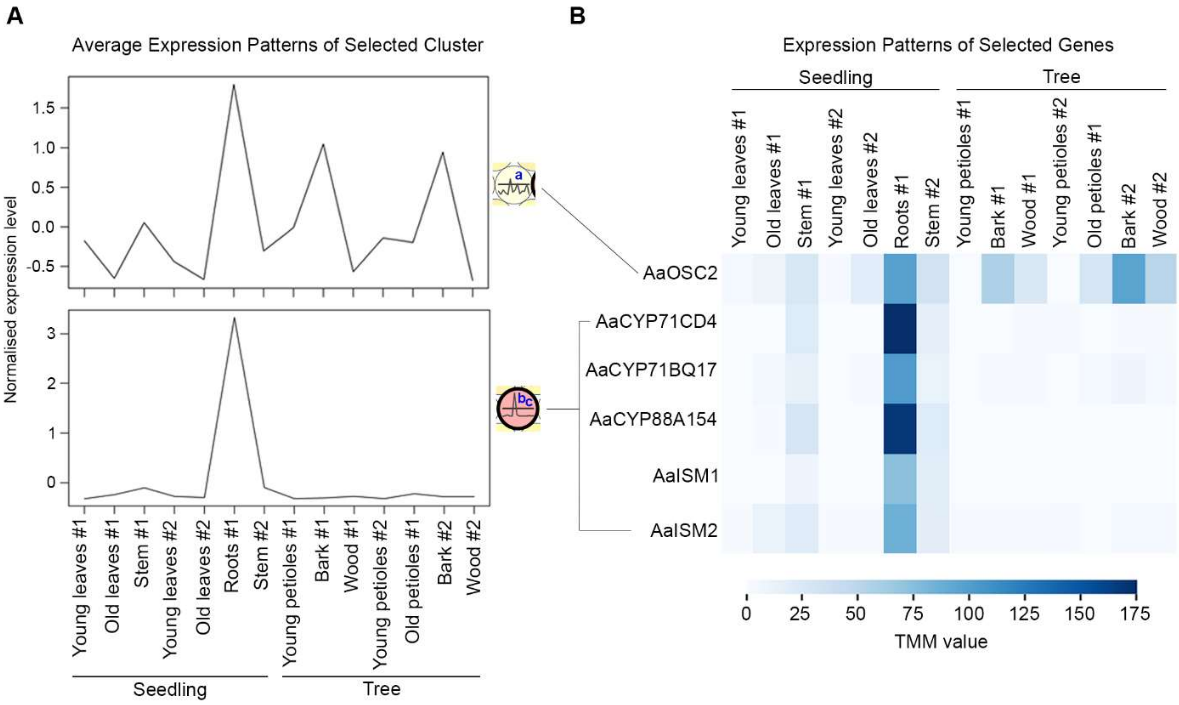


**Figure S1. Comparison of average expression patterns of the two clusters selected from SOM analysis to pathway gene expression patterns.** (A) Average expression pattern of the cluster containing *AaOSC2* (199 contigs in total) (top), and of the cluster containing *AaCYP71CD4* and *AaCYP71BQ17* (695 contigs in total). (B) Expression patterns of verified quassinoid pathway genes as a heat map. The expression level is represented by trimmed mean of M-values (TMM) values in the different tissues of *Ailanthus altissima.* Lines between (A) and (B) connect the genes with the corresponding clusters from SOM analysis.

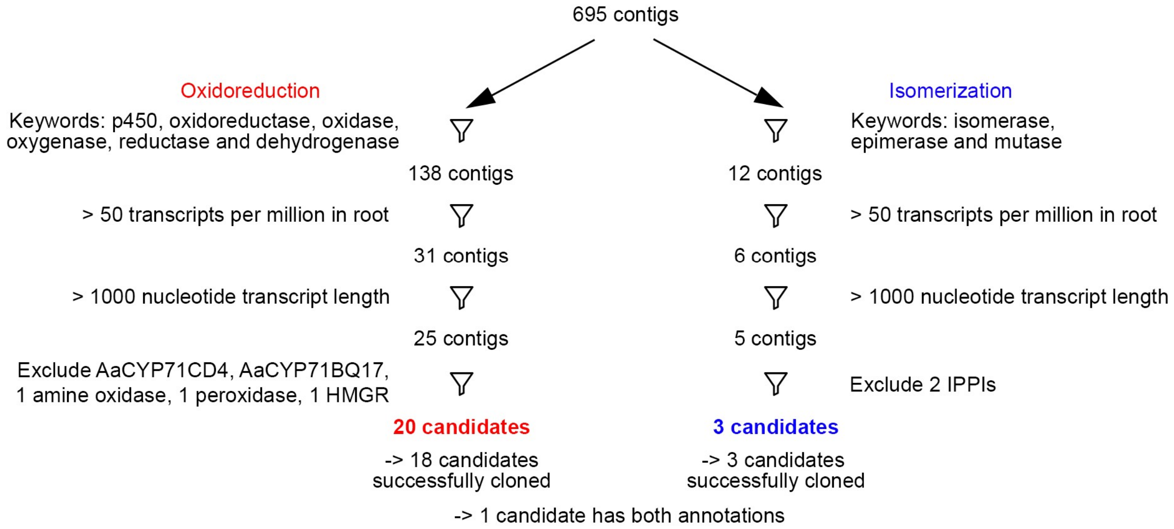


**Figure S2. Candidate selection from self-organising map (SOM) analysis of *Ailanthus altissima* transcriptome data.** The 695 contigs found in the cluster with *AaCYP71CD4* and *AaCYP71BQ17* were first filtered by keywords in their blastn, blastx and pfam annotations. This results in 138 contigs for contigs with oxidoreduction annotations and 12 contigs with isomerization annotations. Among them, 5 contigs were found with both annotations. Contigs with low expression level in root and insufficient nucleotide length were then excluded. Also, candidates with unrelated functions based on their full annotations as well as the two reference CYP450s were manually removed. The resulting contigs were selected as candidates. Abbreviations: isopentenyl-diphosphate delta isomerase (IPPI) and 3-hydroxy-3-methylglutaryl-CoA reductase (HMGR).

**Table S1. Sequences of the primers used in this study.** The start and stop codons are marked in red. For In-Fusion cloning, the overlapping sequences to the vector are marked in blue. For Golden gate cloning, BsaI restriction sites are marked in green, and the cutting site is indicated with a slash

| Primer name | Primer sequence | Vector |
|---|---|---|
| AaOSC2_Fw | CACCACAGGTCTCG/AAAAATGTGGAGGCTTAAGATTGCAGA | pHREAC |
| AaOSC2_Rv | CACCACAGGTCTCG/AGCGTCAATTAGGCAATGGAACTTTCCT | pHREAC |
| AaCYP71CD4_Fw | GCCCAAATTCGCGACCGGATGATGGAGCTACAGCTTGA | pEAQ-HT |
| AaCYP71CD4_Rv | CAGAGTTAAAGGCCTCGATCACGGATCGTAAGGAGTGG | pEAQ-HT |
| AaCYP71BQ17_Fw | GCCCAAATTCGCGACCGGTTGAGAACAAAATTGCCAATGGA | pEAQ-HT |
| AaCYP71BQ17_Rv | CAGAGTTAAAGGCCTCGATCACTTCTGGAAAGGAATATGAGTG | pEAQ-HT |
| AaCYP88A154_Fw | GCCCAAATTCGCGACCGGATGCTAAGAAACTCAGACATCA | pEAQ-HT |
| AaCYP88A154_Rv | CAGAGTTAAAGGCCTCGATCATTTGAGCTTAACGACTTTTGC | pEAQ-HT |
| AaISM2_Fw | CACCACAGGTCTCG/AAAAATGAGCAACCATCCCTATTCTCC | pHREAC |
| AaISM2_Rv | CACCACAGGTCTCG/AGCGTCAGTAGAATTTGGCTTTCTTCTGA | pHREAC |
| AaISM1_Fw | CACCACAGGTCTCG/AAAAATGAGCAATTCGTATATGCCCA | pHREAC |
| AaISM1_Rv | CACCACAGGTCTCG/AGCGTCAGCAAACTTTGGCCTTCT | pHREAC |

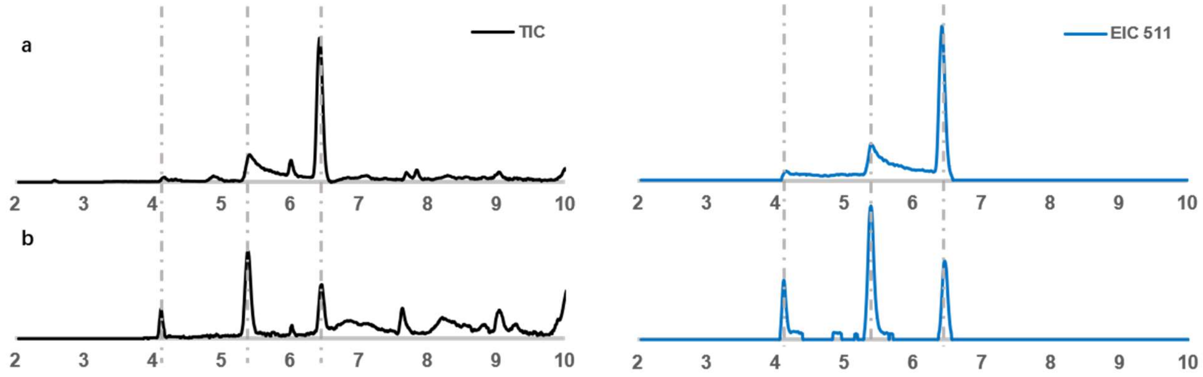


**Figure S3. Acid sensitivity of 7,8-epoxymelianol (9) to silica.** Sample composition of the crude reaction product from melianol epoxidation with *m*-CPBA before (a) and after (b) attempts to purify the 6.6 min peak corresponding to 7,8-epoxymelianol (**9**) by normal phase column chromatography on silica. Data shown are LCMS chromatograms (black: TIC; blue: EIC 511).

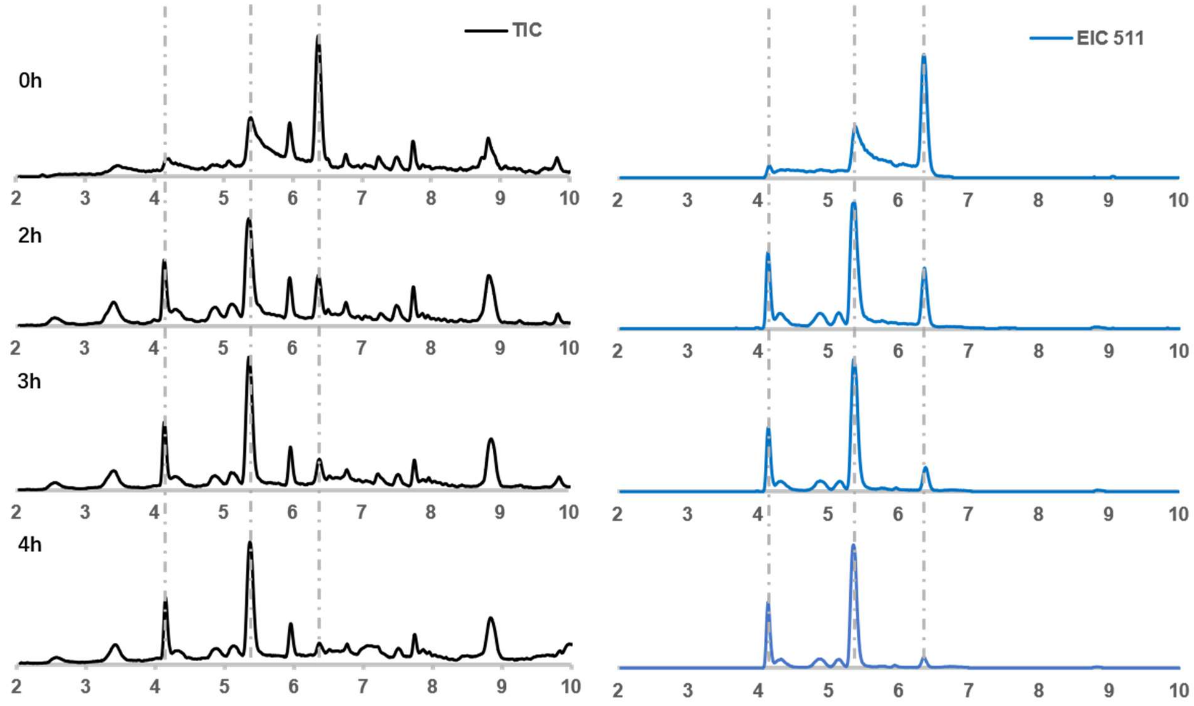


**Figure S4. Time course showing degradation of 7,8-epoxymelianol (9) (peak at 6.6 min) in 50/50 MeCN/$H_2O$ + 0.1% formic acid.** Data shown are LCMS chromatograms (black: TIC; blue: EIC 511).

**Table S2. Conditions for extraction and flash chromatography purification of isomeliandiol (10) and protoglabretal (11).**

| | | | | |
|---|---|---|---|---|
| **Isomeliandiol (10)** | **Fresh weight:** | 82.27 g | **Dry weight:** | 16.06 g |
| | **Extraction solvent:** | Ethyl acetate | **Crude extract:** | 1.18 g |
| | **Column** | **Solvents** | **Gradient** | **Yield** |
| | SNAP KP-Sil 50 g | A: Petroleum ether<br>B: Ethyl acetate | 10-100% (10 CV)<br>100% (3 CV) | 164 mg |
| | Sfär C18 D 12 g | A: Water<br>B: Acetonitrile | 40-100% (11 CV) | 27 mg |
| | SNAP Ultra 10 g | A: Petroleum ether<br>B: Ethyl acetate | 30% (2 CV)<br>30-60% (3 CV)<br>60% (7 CV)<br>60-80% (3 CV)<br>80% (3 CV)<br>80%-100 (3 CV) | 9 mg |
| | Sfär C18 D 12 g | A: Water<br>B: Acetonitrile | 60-100% (11 CV) | 2 mg |
| **Protoglabretal (11)** | **Fresh weight:** | 114.95 g | **Dry weight:** | 17.59 g |
| | **Extraction solvent:** | 90/10 MeOH/$H_2O$ | **Crude extract:** | 5.77 g |
| | **Column** | **Solvents** | **Gradient** | **Yield** |
| | SNAP KP-Sil 100 g | A: Petroleum ether<br>B: Ethyl acetate | 10-100% (10 CV)<br>100% (3 CV) | 140 mg |
| | Sfär C18 D 12 g | A: Water<br>B: Acetonitrile | 60-100% (11 CV) | 22 mg |
| | SNAP Ultra 10 g | A: Petroleum ether<br>B: Ethyl acetate | 30% (2 CV)<br>30-67% (3 CV)<br>67% (5 CV)<br>67-100% (3 CV)<br>100% (5 CV) | 12 mg |

**Table S3.** $^{1}H$ **and** $^{13}C$ **NMR data of 7,8-epoxymelianol (9) ($C_6D_6$, 298 K, 600 MHz).**

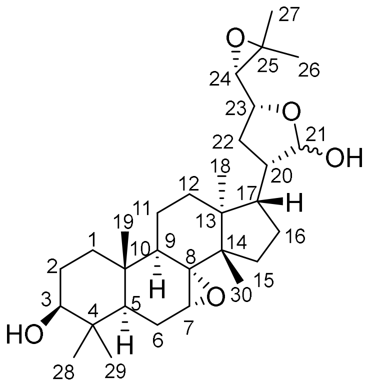


| Atom | $^{13}C$ ppm [a] | $^{1}H$ ppm (m, Hz) [a] |
|---|---|---|
| 1 | 38.05 / 37.98 | 0.77-0.83, 1.34-1.46 (m, 2H) |
| 2 | 27.87 | 1.31-1.38 (m, 2H) |
| 3 | 78.47 / 78.40 | 2.95-2.99 (m, 1H) |
| 4 | 38.60 | - |
| 5 | 45.45 / 45.37 | 1.25-1.30 (m, 1H) |
| 6 | 23.21 / 23.20 | 1.40-1.46, 1.91-1.96 (m, 2H) |
| 7 | 55.05 / 54.96 | 2.70 (t, *J* = 1.7 Hz, 1H) |
| 8 | 63.22 / 63.10 | - |
| 9 | 49.89 / 49.80[b] | 1.85-1.91 (m, 1H) |
| 10 | 35.08 | - |
| 11 | 17.65 / 17.61 | 1.42-1.48, 1.52-1.58 (m, 2H) |
| 12 | 31.18 / 31.52 | 1.41-1.46, 1.98-2.04 (m, 2H) / 1.57-1.61, 1.78-1.93 (m, 2H) |
| 13 | 45.34 / 45.61 | - |
| 14 | 49.09 / 49.38 | - |
| 15 | 28.29 / 27.84 | 0.82-0.87, 1.98-2.05 (m, 2H) / 0.80-0.82, 1.33-1.45 (m, 2H) |
| 16 | 26.94 / 27.08 | 1.20-1.26, 1.71-1.81 (m, 2H) |
| 17 | 45.89 / 51.25 | 2.17-2.20 (m, 1H) / 1.70-1.73 (m, 1H) |
| 18 | 22.02 / 21.24 | 1.09 (s, 3H) /1.28 (s, 3H) |
| 19 | 14.53 / 14.55 | 0.67 (s, 3H) / 0.70 (s, 3H) |
| 20 | 47.19 / 49.72[b] | 1.81-1.87 (m, 1H) / 2.17-2.21 (m, 1H) |
| 21 | 97.87 / 102.2 | 5.29 (d, *J* = 3.9 Hz, 1H) / 5.31 (d, *J* = 3.4 Hz, 1H) |
| 22 | 32.05 / 35.78 | 1.61-1.72 (m, 2H) / 1.15-1.20, 1.80-1.84 (m, 2H) |
| 23 | 78.59 / 77.43 | 3.87-3.93 (m, 1H) / 4.02 (ddd, *J* = 10.7, 7.3, 5.1 Hz, 1H) |
| 24 | 67.89 / 65.52 | 2.91 (d, *J* = 7.2 Hz, 1H) / 2.78 (d, *J* = 7.3 Hz, 1H) |
| 25 | 57.41 / 56.41 | - |
| 26 | 25.17 / 25.02 | 1.14 (s, 3H) / 1.12 (s, 3H) |
| 27 | 19.32 / 19.59 | 1.12 (s, 3H) /1.09 (s, 3H) |
| 28 | 28.09 / 28.05 | 0.97 (s, 3H) |
| 29 | 16.04 / 16.06 | 0.84 (s, 3H) |
| 30 | 22.97 / 22.81 | 0.94 (s, 3H) / 0.90 (s, 3H) |

[a] Second value corresponds to the minor lactol epimer whenever clearly discernible.

[b] Signals interchangeable.

**Table S4. $^1$H and $^{13}$C NMR data of isomeliandiol (10) obtained as a mixture of epimers ($CDCl_3$, 298 K, 500 MHz).**

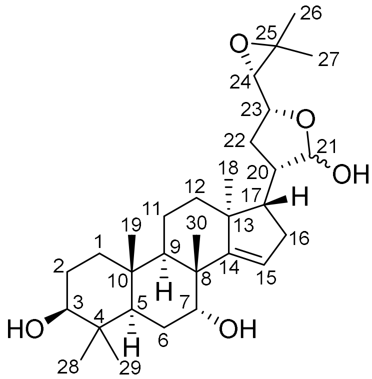

| Atom | $^{13}$C ppm [a] | $^1$H ppm (m, Hz) [a] |
|---|---|---|
| 1 | 38.05 / 38.03 | 1.03-1.07, 1.57-1.63 (m, 2H) |
| 2 | 27.29 | 1.55-1.66 (m, 2H) |
| 3 | 78.92 / 78.89 | 3.27 (dd, *J* = 11.1 Hz, 1H) / 3.28 (dd, *J* = 4.7 Hz, 1H) |
| 4 | 38.51 | - |
| 5 | 46.68 / 46.64 | 1.45-1.52 (m, 1H) |
| 6 | 23.81 / 23.83 | 1.70-1.78, 1.82-1.88 (m, 2H) |
| 7 | 72.47 / 72.48 | 3.92 (m, 1H) |
| 8 | 44.39 / 44.38 | - |
| 9 | 41.88 / 41.93 | 1.88-1.94 (m, 1H) |
| 10 | 37.73 | - |
| 11 | 16.48 / 16.47 | 1.48-1.53, 1.67-1.72 (m, 2H) |
| 12 | 33.22 / 33.27 | 1.49-1.53, 1.77-1.81 (m, 2H) |
| 13 | 46.85 / 46.78 | - |
| 14 | 162.35 / 162.22 | - |
| 15 | 119.67 / 119.18 | 5.47 (m, 1H) / 5.45 (m, 1H) |
| 16 | 35.22 / 35.14 | 2.15-2.20 (m, 2H) |
| 17 | 52.88 / 52.96 | 1.92-2.01 (m, 1H) |
| 18 | 19.94 / 19.96 | 1.02 (s, 3H) |
| 19 | 15.54 | 0.89 (s, 3H) |
| 20 | 45.63 / 47.95 | 1.46-1.51 (m, 1H) |
| 21 | 97.74 / 102.55 | 5.38 (d, *J* = 3.2 Hz, 1H) |
| 22 | 31.50 / 34.87 | 1.69-1.77, 2.00-2.05 / 1.36-1.44, 2.07-2.15 (m, 2H) |
| 23 | 78.53 / 77.36 | 3.88-3.94 (m, 1H) / 3.93-3.99 (m, 1H) |
| 24 | 67.77 / 65.35 | 2.82 (d, *J* = 7.4 Hz, 1H) / 2.69 (d, *J* = 7.6 Hz, 1H) |
| 25 | 58.27 / 57.46 | - |
| 26 | 25.17 / 25.06 | 1.32 (s, 3H) / 1.33 (s, 3H) |
| 27 | 19.35 / 19.59 | 1.32 (s, 3H) / 1.09 (s, 3H) |
| 28 | 27.81 | 0.99 (s, 3H) |
| 29 | 15.59 | 0.79 (s, 3H) |
| 30 | 27.80 | 1.05 (s, 3H) |

[a] Second value corresponds to the minor lactol epimer whenever clearly discernible.

**Table S5. Partial $^{1}$H and $^{13}$C NMR data of degradation products 12 and 13 present in NMR samples of isomeliandiol (10) ($CDCl_3$, 298 K, 500 MHz).**

| | 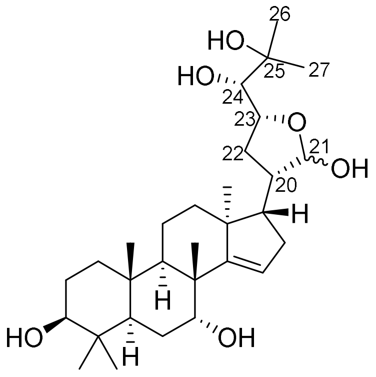 **12** | | 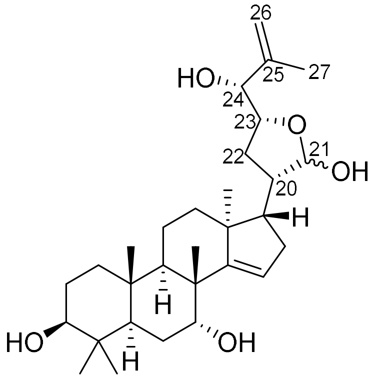 **13** | |
|---|---|---|---|---|
| **Atom** | **$^{13}$C ppm** | **$^{1}$H ppm** | **$^{13}$C ppm** | **$^{1}$H ppm** |
| 20 | 45.13 | 2.17 | 45.79 | 2.22 |
| 21 | 97.54 | 5.33 | 97.30 | 5.29 |
| 22 | 30.19 | 1.95 | 30.52 | 1.81 |
| 23 | 78.69 | 4.46 | 80.31 | 4.23 |
| 24 | 75.37 | 3.19 | 77.23 | 3.90 |
| 25 | 73.35 | - | 145.09 | - |
| 26 | 26.77 | 1.29 | 112.62 | 4.94, 5.04 |
| 27 | 26.91 | 1.27 | 18.87 | 1.78 |

**Table S6.** $^1$H and $^{13}$C NMR data of protoglabretal (11) ($CDCl_3$, 298 K, 500 MHz).

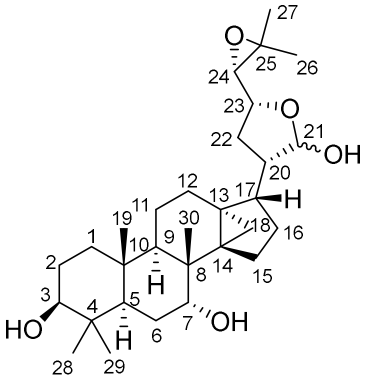


| Atom | $^{13}$C ppm [a] | $^1$H ppm (m, Hz) [a] |
|---|---|---|
| 1 | 38.70 / 38.58 | 0.92-1.00, 1.57-1.63 (m, 2H) |
| 2 | 27.28 | 1.59-1.67 (m, 2H) |
| 3 | 78.92 / 78.89 | 3.26 (dd, *J* = 11.4, 4.3 Hz, 1H) |
| 4 | 38.47 / 38.46 | - |
| 5 | 46.11 / 46.03 | 1.43-1.48 (m, 1H) |
| 6 | 24.35 / 34.34 | 1.56-1.63, 1.71-1.77 (m, 2H) |
| 7 | 74.65 / 74.51 | 3.74-3.79 (m, 1H) |
| 8 | 38.90 / 38.97 | - |
| 9 | 44.32 / 44.12 | 1.16-1.24 (m, 1H) |
| 10 | 37.34 / 37.40 | - |
| 11 | 25.75 | 1.70-1.78, 2.06-2.14 (m, 2H) |
| 12 | 16.46 / 16.30 | 1.23-1.36 (m, 2H) |
| 13 | 29.07 / 28.72 | - |
| 14 | 37.03 / 36.23 | - |
| 15 | 26.45 / 27.26 | 1.88-1.95 (m, 2H) / 1.49-1.58 (m, 2H) |
| 16 | 27.63 / 26.29 | 0.85-0.88, 1.62-1.68 (m, 2H) / 0.89-0.93, 1.62-1.68 (m, 2H) |
| 17 | 44.90 / 48.44 | 2.15-2.24 (m, 1H) / 1.99-2.05 (m, 1H) |
| 18 | 13.88 / 13.65 | 0.45 (d, *J* = 4.9 Hz, 1H), 0.65 (d, *J* = 4.8 Hz, 1H) / 0.47 (d, *J* = 5.3 Hz, 1H), 0.73 (d, *J* = 5.1 Hz, 1H) |
| 19 | 16.06 / 15.93 | 0.87 (s, 3H) |
| 20 | 49.48 / 50.91 | 1.82-1.90 (m, 1H) / 2.10-2.15 (m, 1H) |
| 21 | 98.34 / 102.25 | 5.42 (d, *J* = 3.9 Hz, 1H) / 5.42 (d, *J* = 3.4 Hz, 1H) |
| 22 | 30.93 / 33.13 | 1.65-1.74, 1.94-1.99 (m, 2H) / 1.37-1.43, 2.02-2.08 (m, 2H) |
| 23 | 78.57 / 77.46 | 3.84-3.91 (m, 1H) /3.92-3.97 (m, 1H) |
| 24 | 67.78 / 65.41 | 2.83 (d, *J* = 7.5 Hz, 1H) /2.69 (d, *J* = 7.6 Hz, 1H) |
| 25 | 58.31 / 57.51 | - |
| 26 | 25.14 / 25.05 | 1.31 (s, 3H) / 1.33 (s, 3H) |
| 27 | 19.32 / 19.56 | 1.30 (s, 3H) / 1.02 (s, 3H) |
| 28 | 27.87 / 27.86 | 0.97 (s, 3H) |
| 29 | 15.68 / 15.67 | 0.77 (s, 3H) |
| 30 | 19.66 / 19.53 | 1.03 (s, 3H) |

[a] Second value corresponds to the minor lactol epimer whenever clearly discernible.

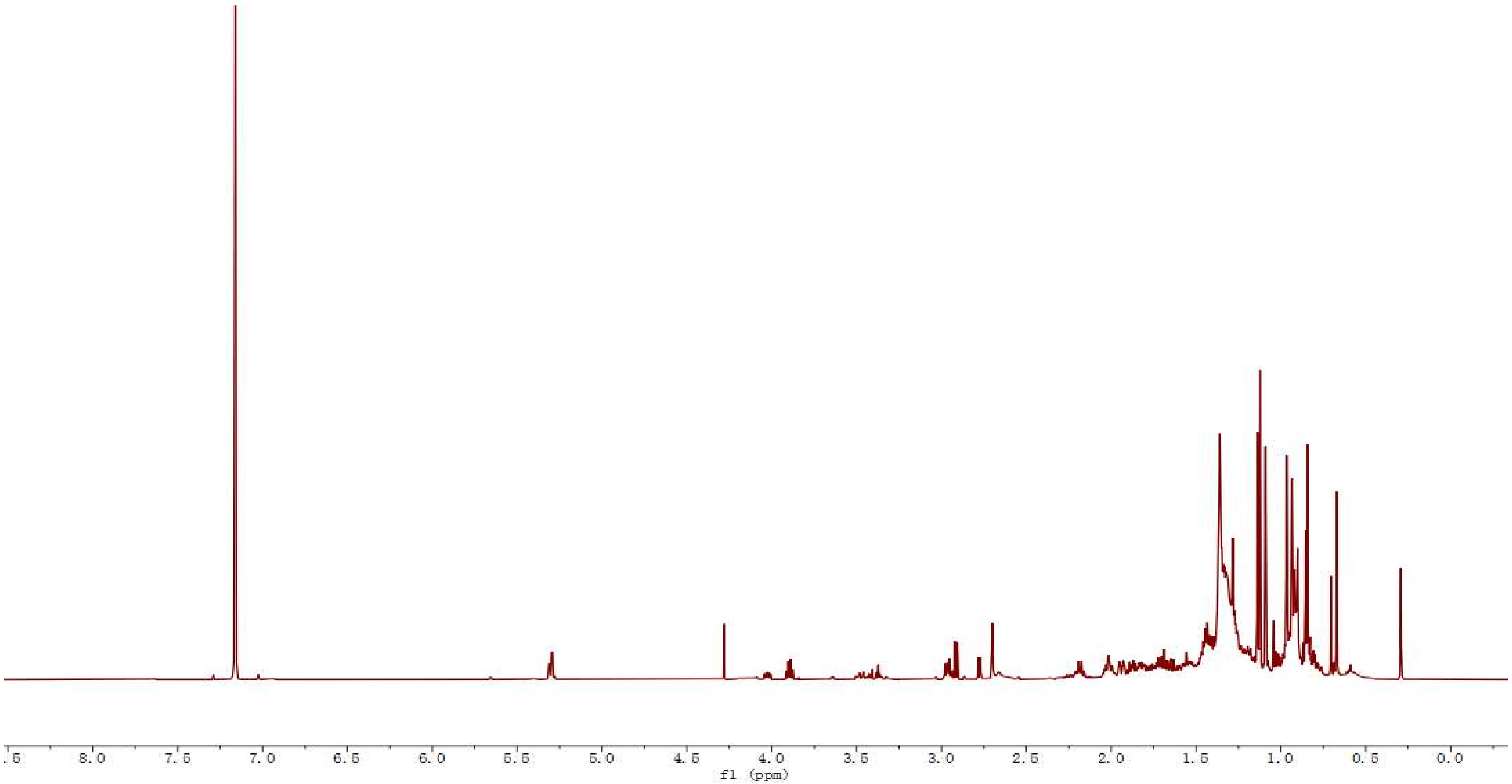


**Figure S5.** [1]**H spectrum of 7,8-epoxymelianol (9) ($C_6D_6$, 298 K, 600 MHz).**

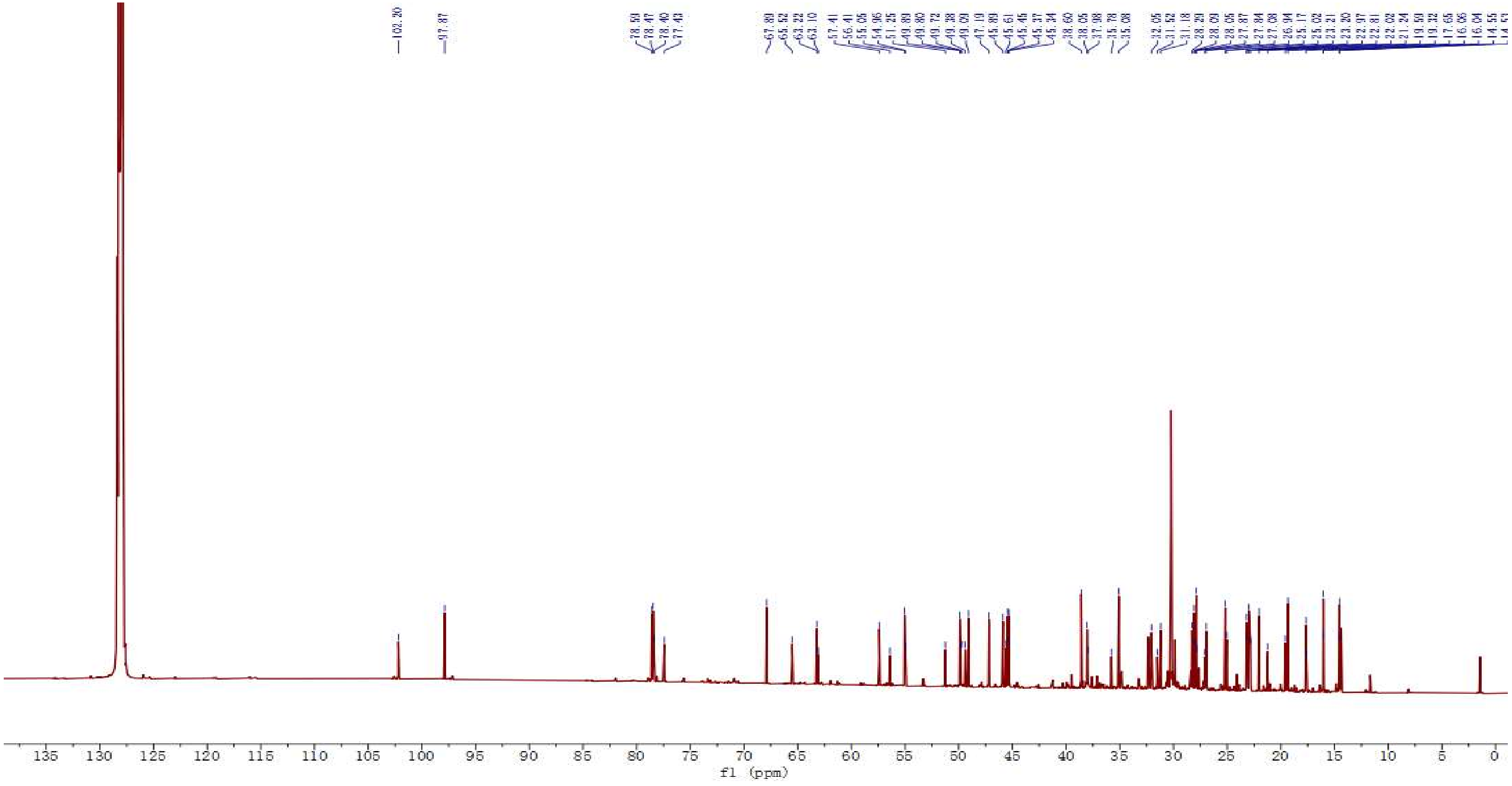


**Figure S6.** ¹³C spectrum of 7,8-epoxymelianol (9) ($C_6D_6$, 298 K, 151 MHz).

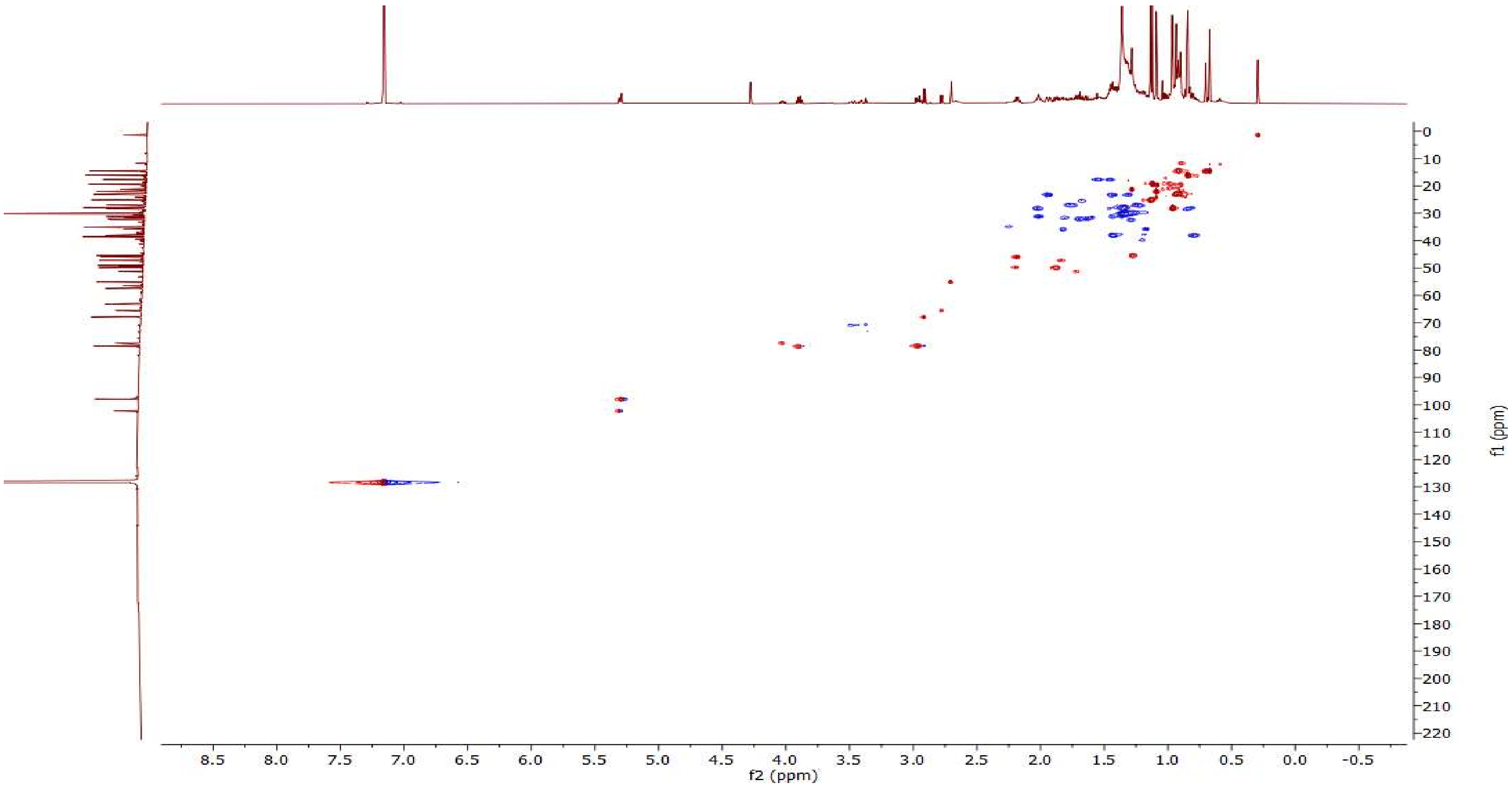


**Figure S7. HSQC spectrum of 7,8-epoxymelianol (9) ($C_6D_6$, 298 K, 600 MHz).**

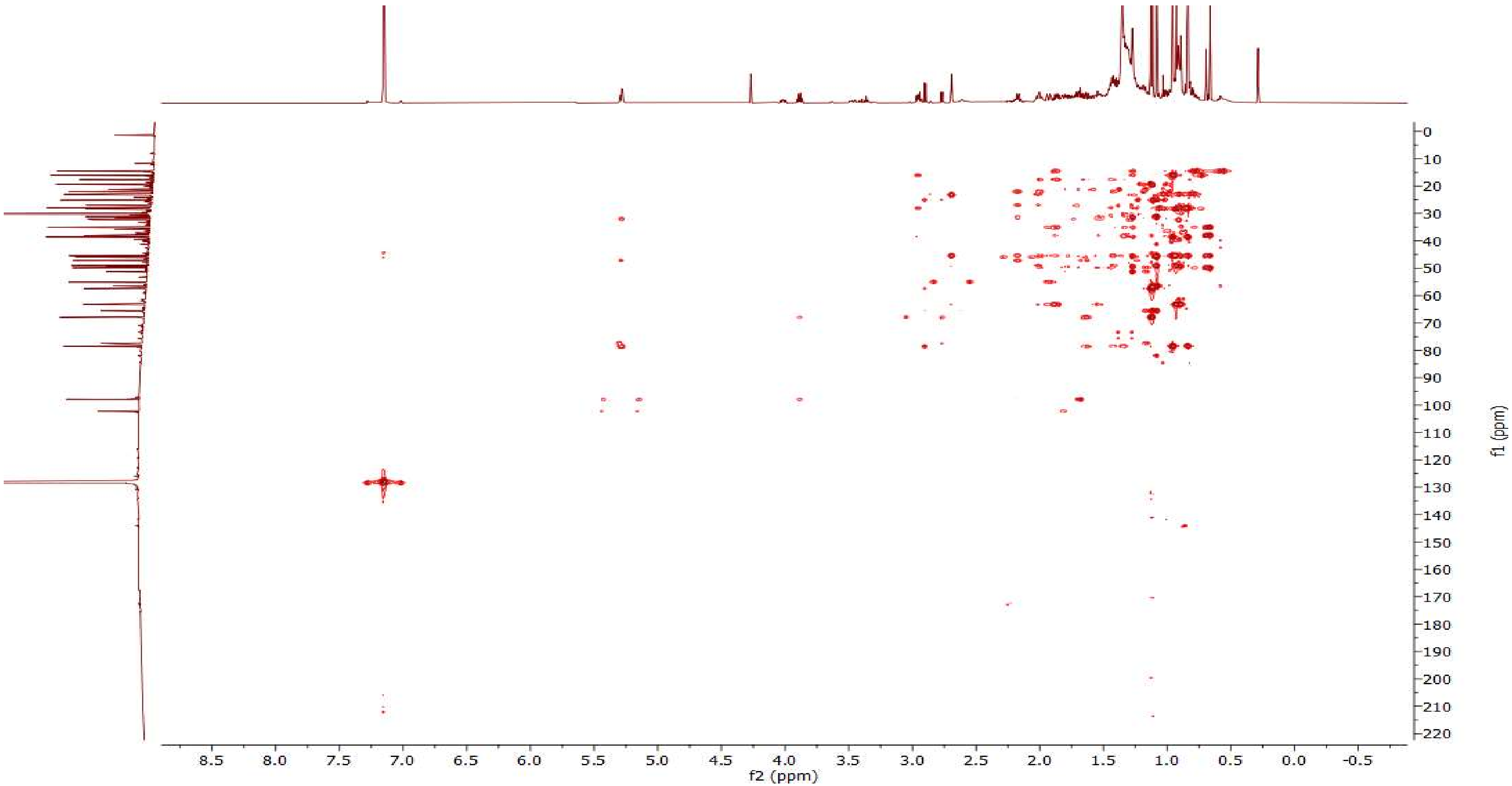


**Figure S8. HMBC spectrum of 7,8-epoxymelianol (9) ($C_6D_6$, 298 K, 600 MHz).**

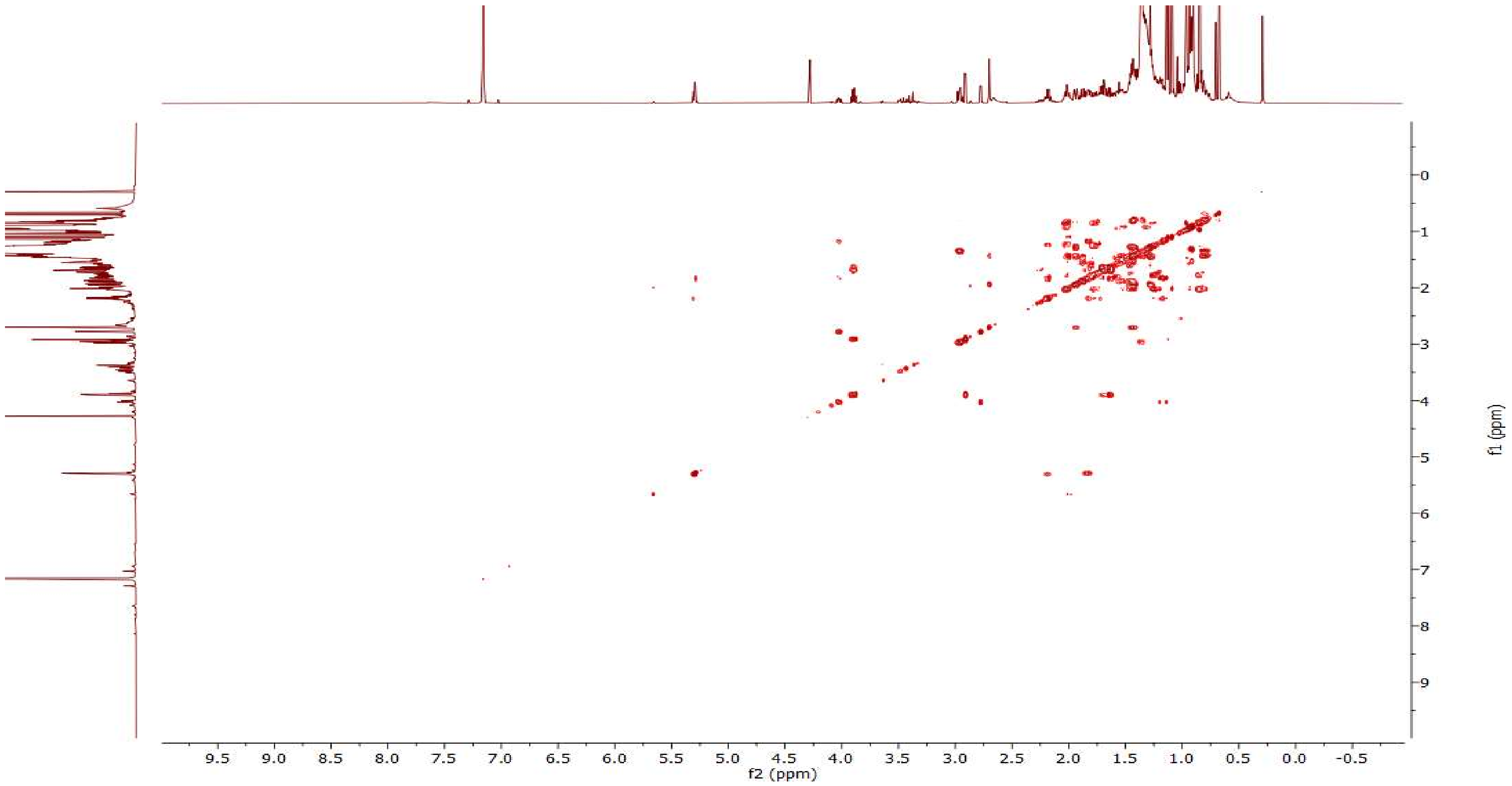


**Figure S9. COSY spectrum of 7,8-epoxymelianol (9) ($C_6D_6$, 298 K, 600 MHz).**

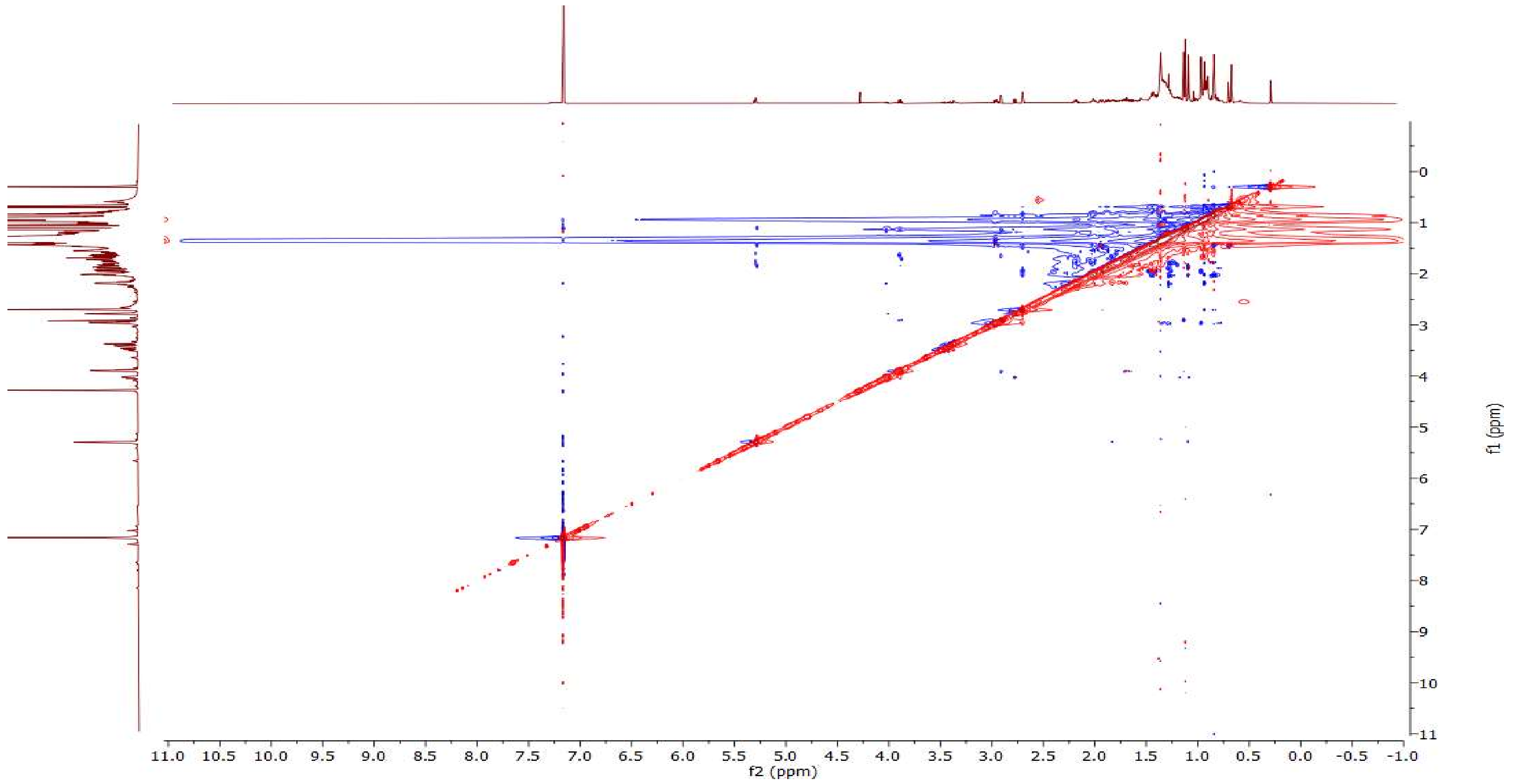


**Figure S10. NOESY spectrum of 7,8-epoxymelianol (9) ($C_6D_6$, 298 K, 500 MHz).**

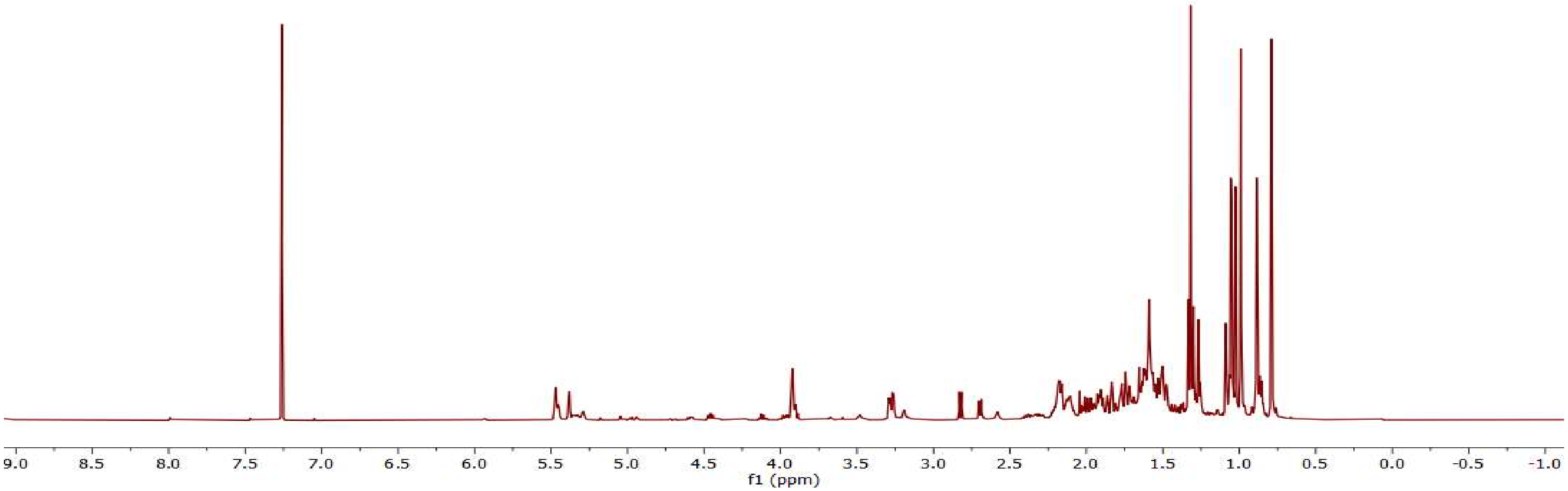


**Figure S11.** **$^1$H NMR spectrum of isomeliandiol (10) ($CDCl_3$, 298 K, 500 MHz).**

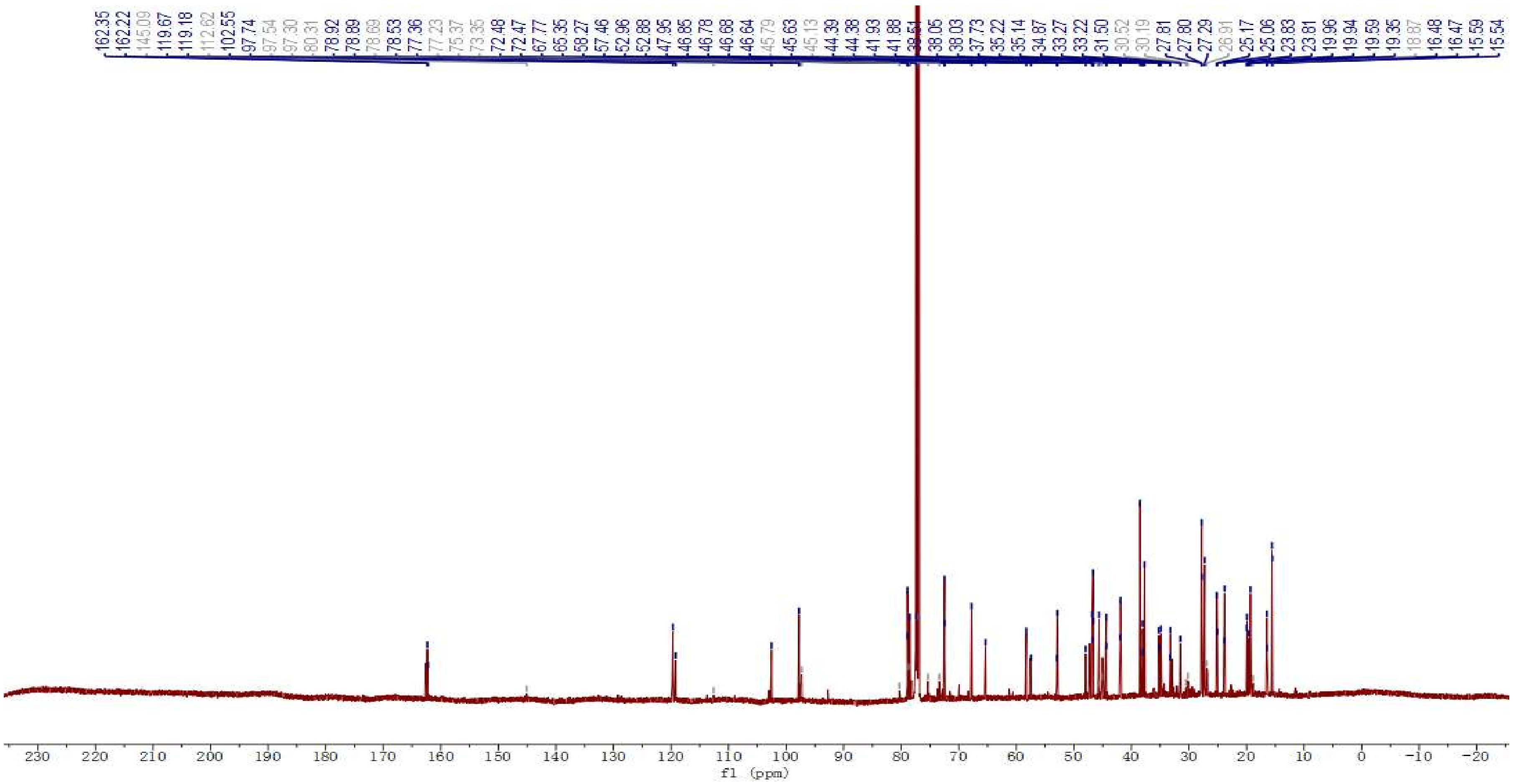


**Figure S12. $^{13}C$ NMR spectrum of isomeliandiol (10) containing the C24/25 epoxide degradation products 12 and 13 ($CDCl_3$, 298 K, 126 MHz).** The $^{13}C$ resonances of isomeliandiol (**10**) are shown in blue and key resonances of degradation products **12** and **13** are shown in grey.

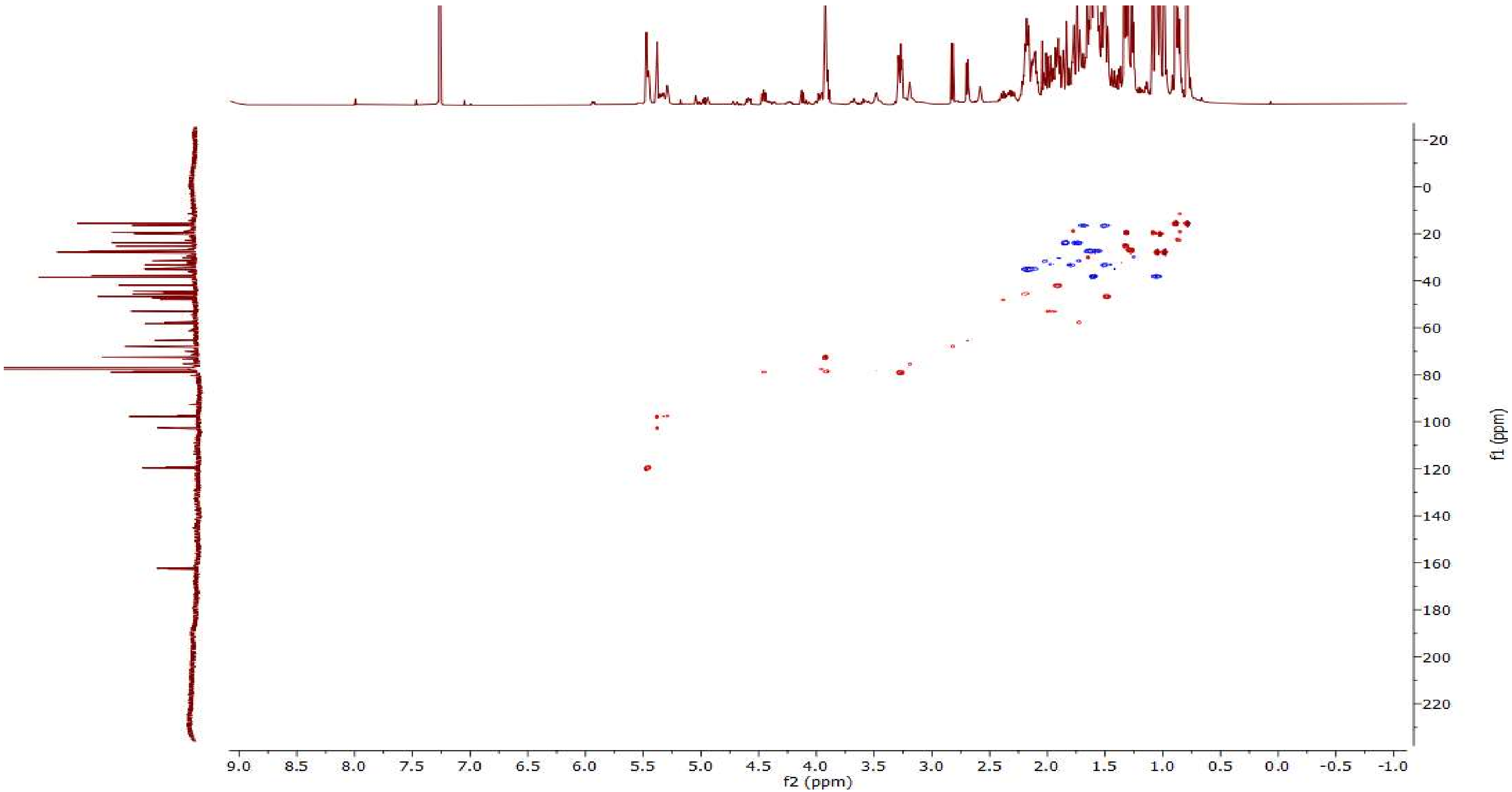


**Figure S13. HSQC spectrum of isomeliandiol (10) ($CDCl_3$, 298 K, 500 MHz).**

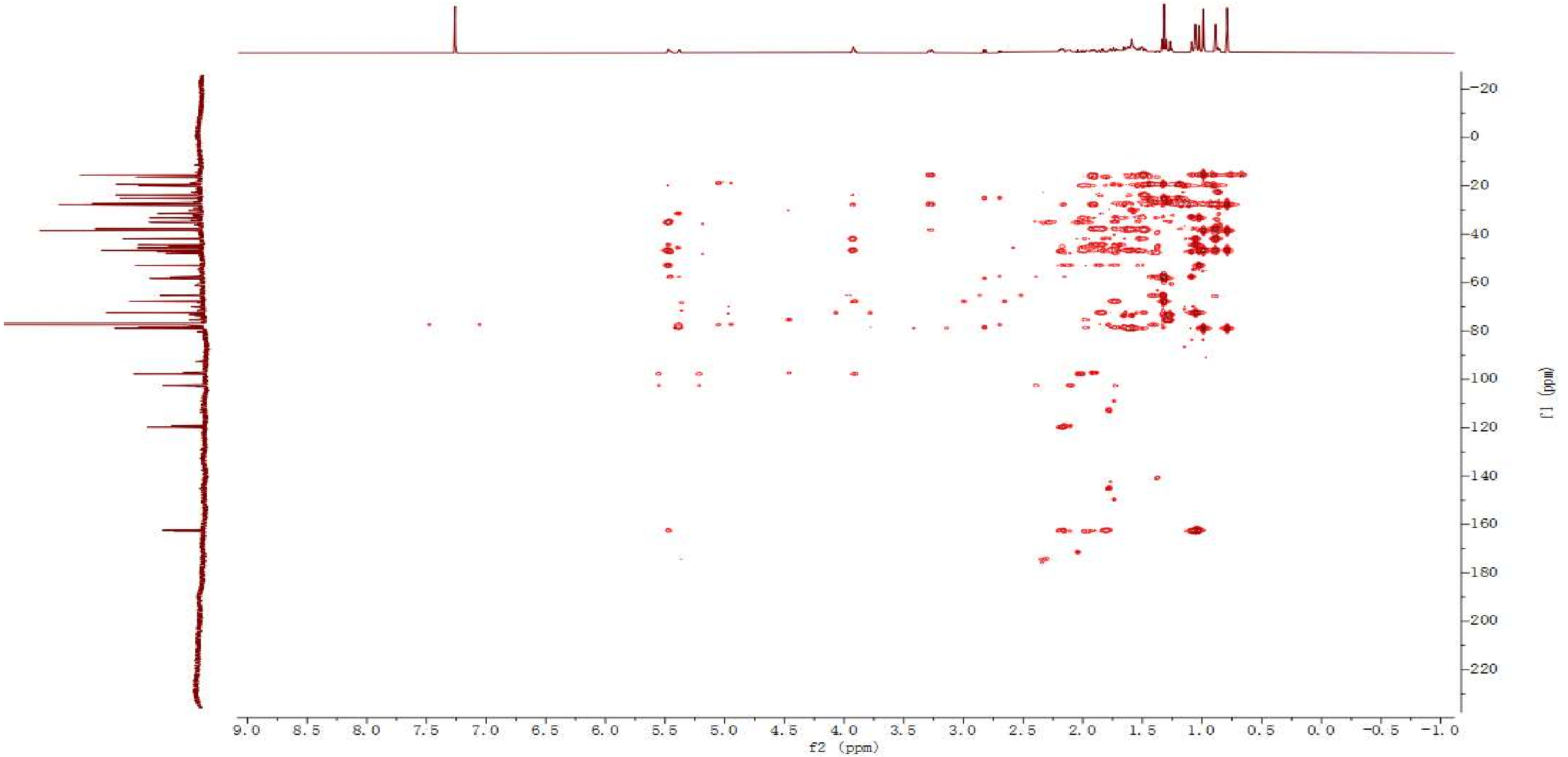


**Figure S14. HMBC spectrum of isomeliandiol (10) ($CDCl_3$, 298 K, 500 MHz).**

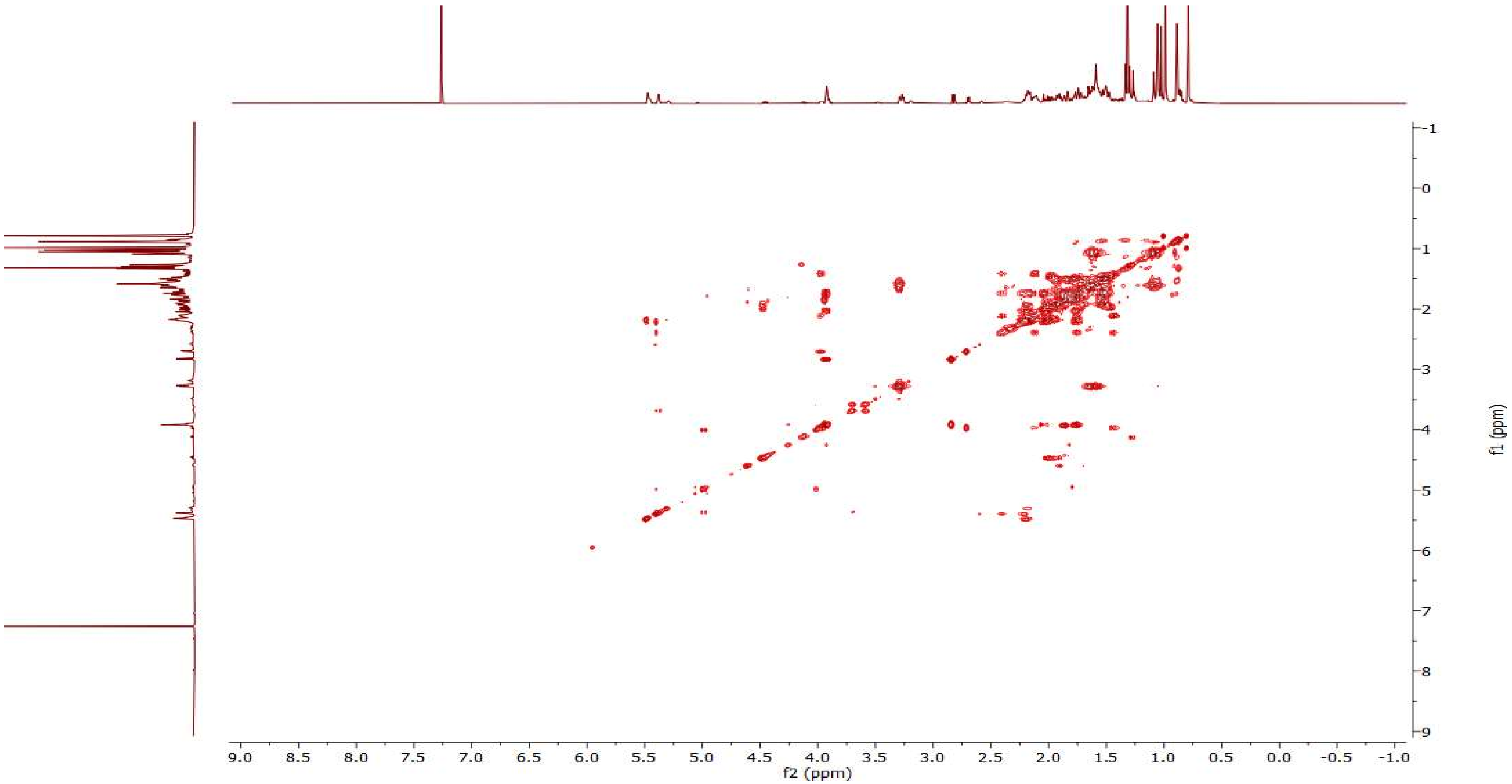


**Figure S15. COSY spectrum of isomeliandiol (10) ($CDCl_3$, 298 K, 500 MHz).**

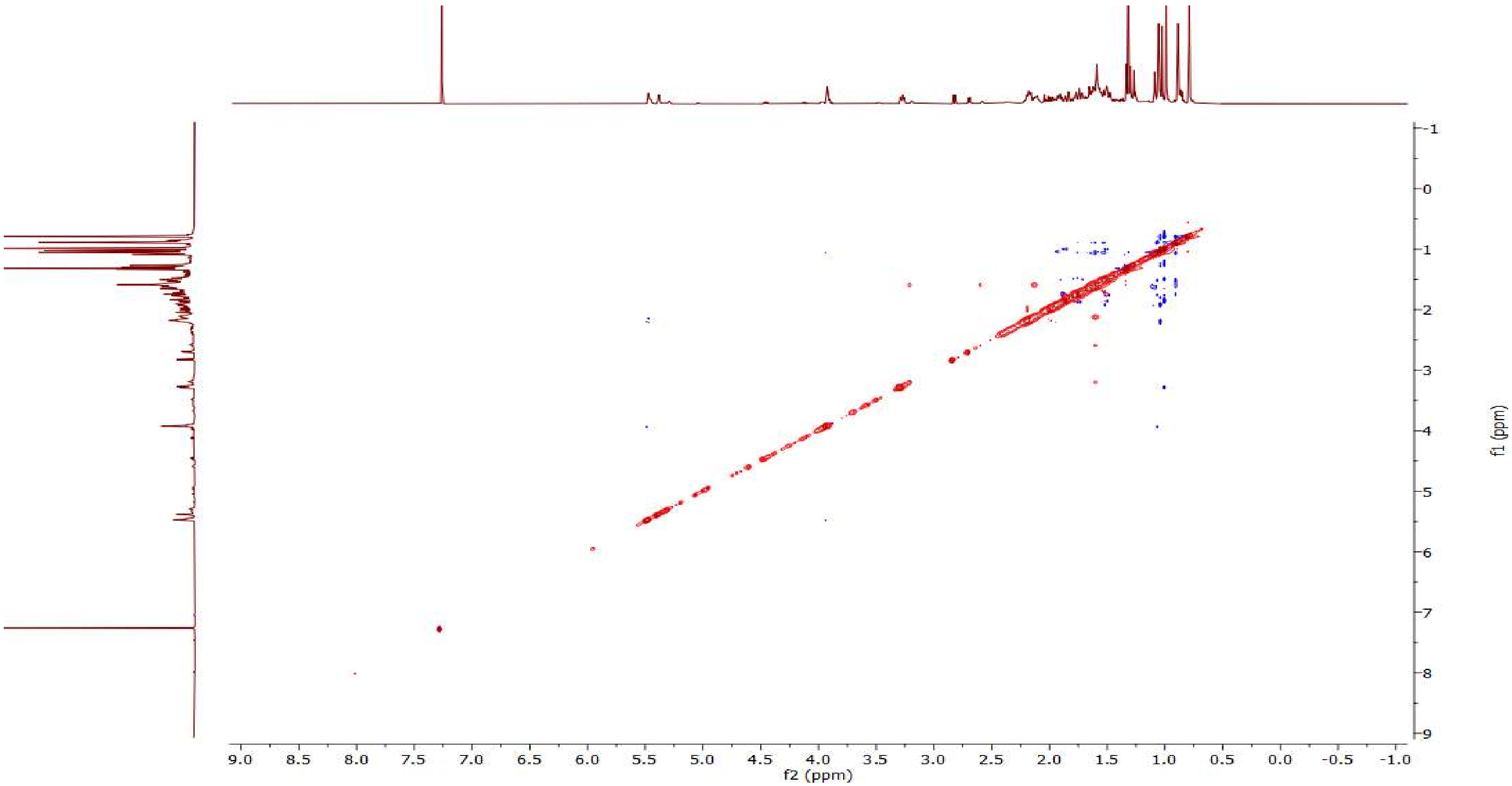


**Figure S16. NOESY spectrum of isomeliandiol (10) ($CDCl_3$, 298 K, 500 MHz).**

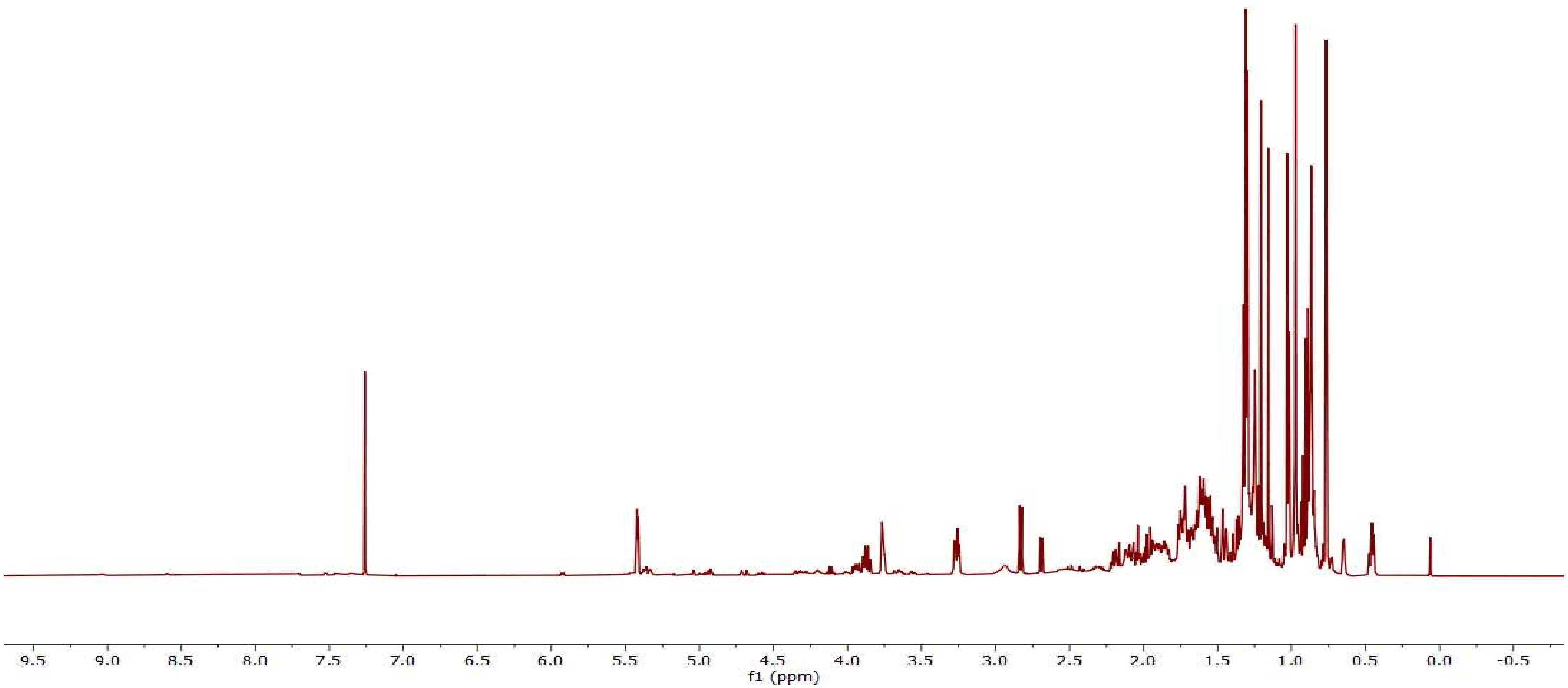


**Figure S17. $^{1}H$ NMR spectrum of protoglabretal (11) ($CDCl_3$, 298 K, 500 MHz).**

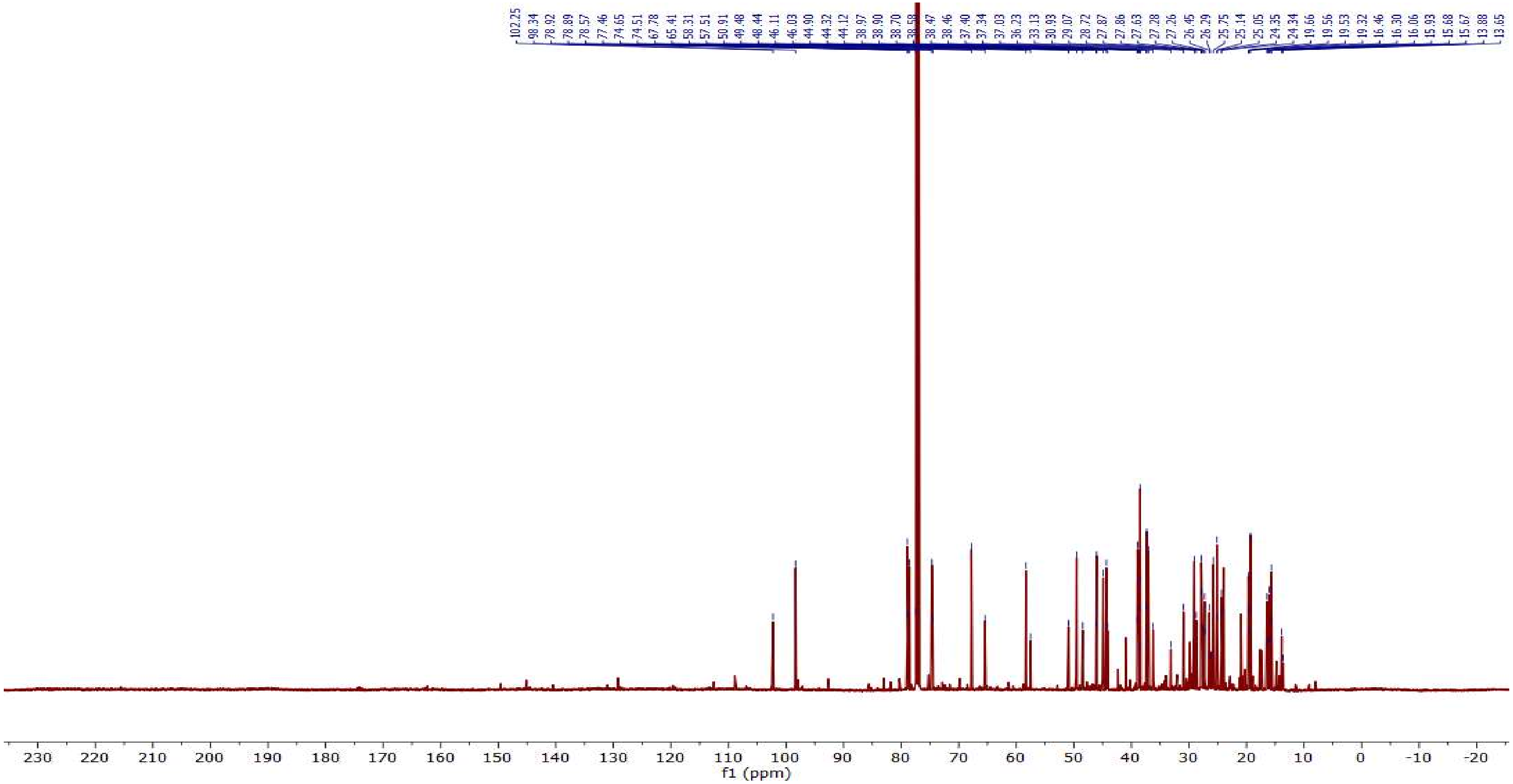


**Figure S18.** **$^{13}C$ NMR spectrum of protoglabretal (11) ($CDCl_3$, 298 K, 126 MHz).**

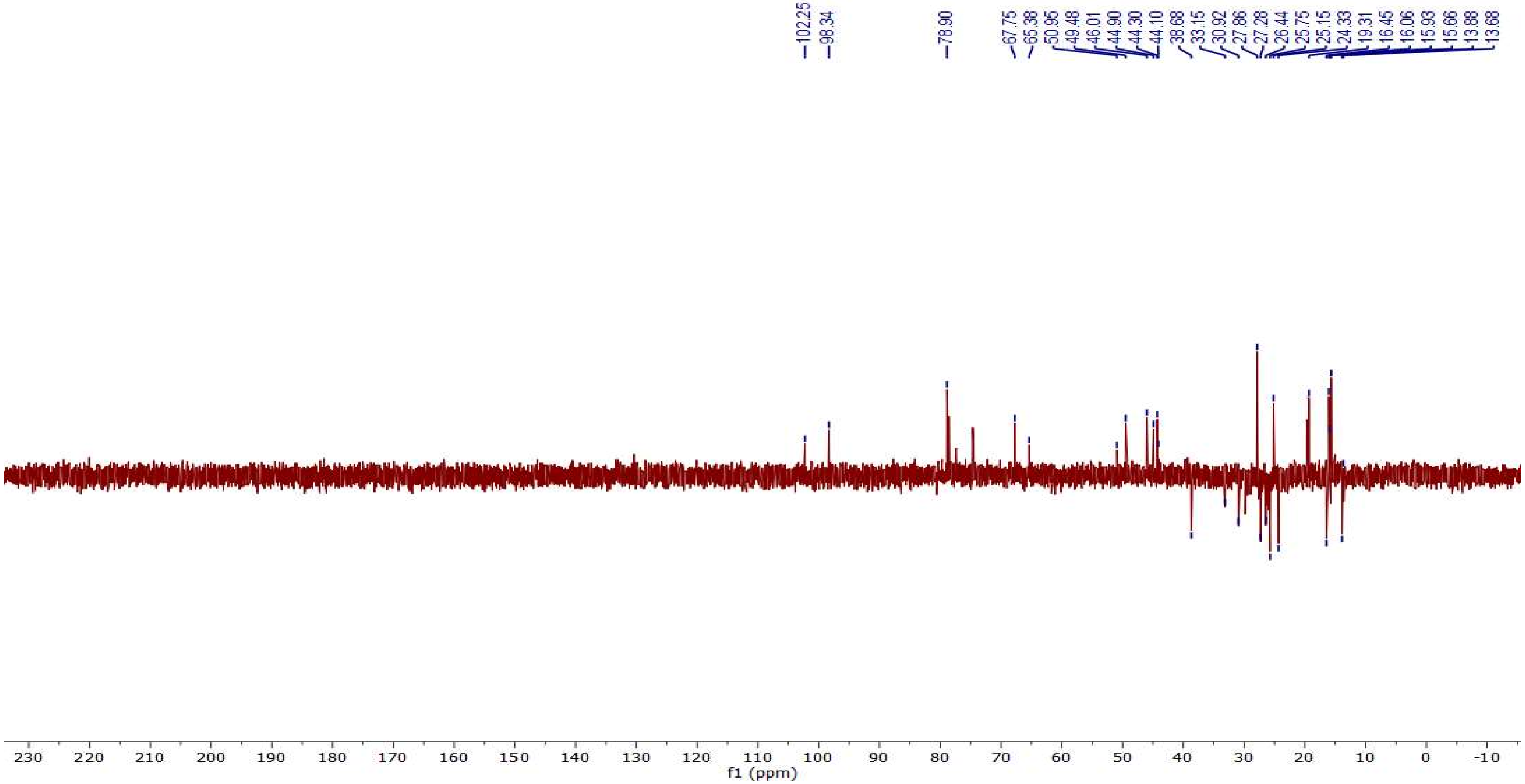


**Figure S19. DEPT-135 spectrum of protoglabretal (11) ($CDCl_3$, 298 K, 100 MHz).**

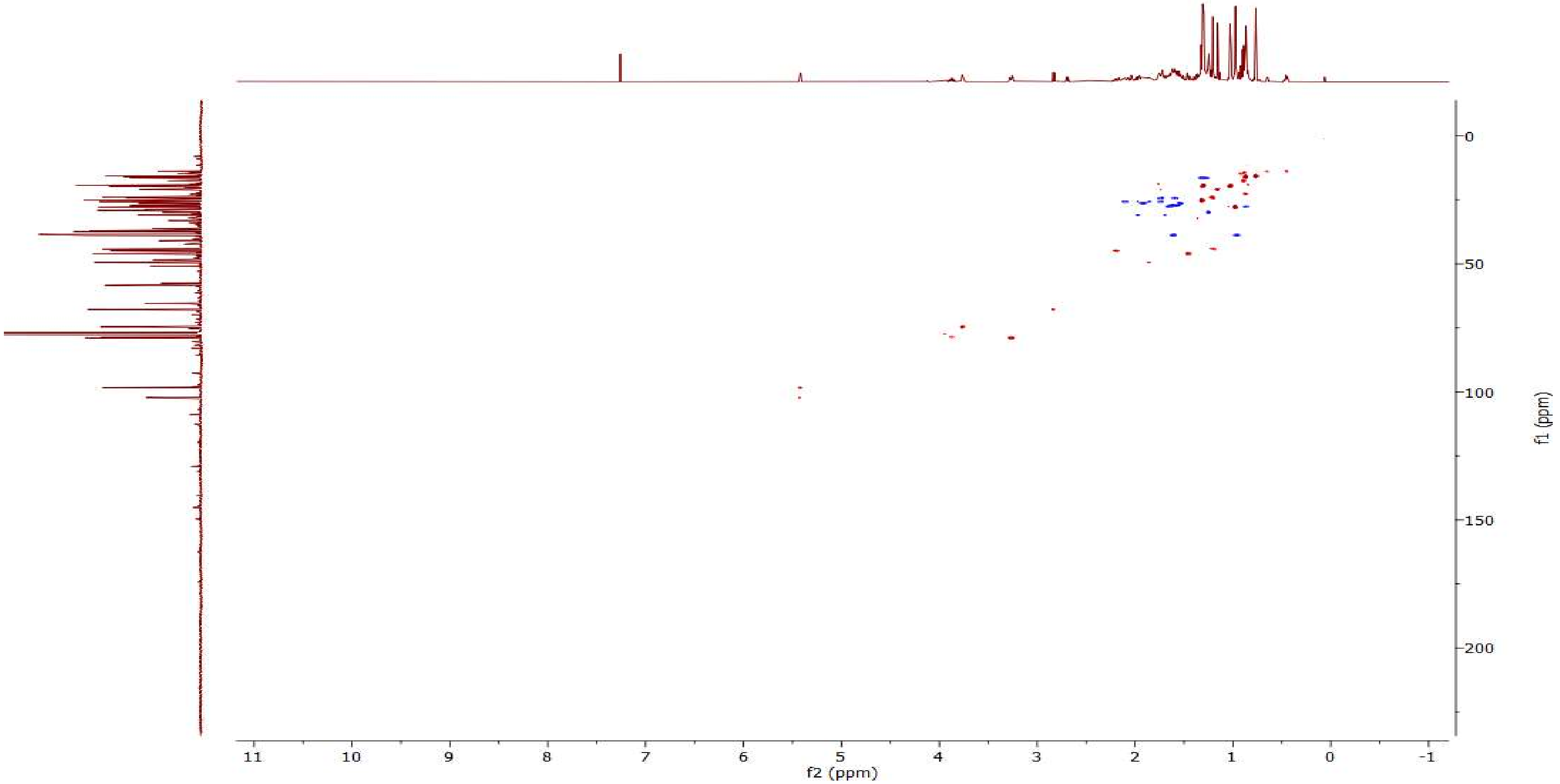


**Figure S20. HSQC spectrum of protoglabretal (11) ($CDCl_3$, 298 K, 500 MHz).**

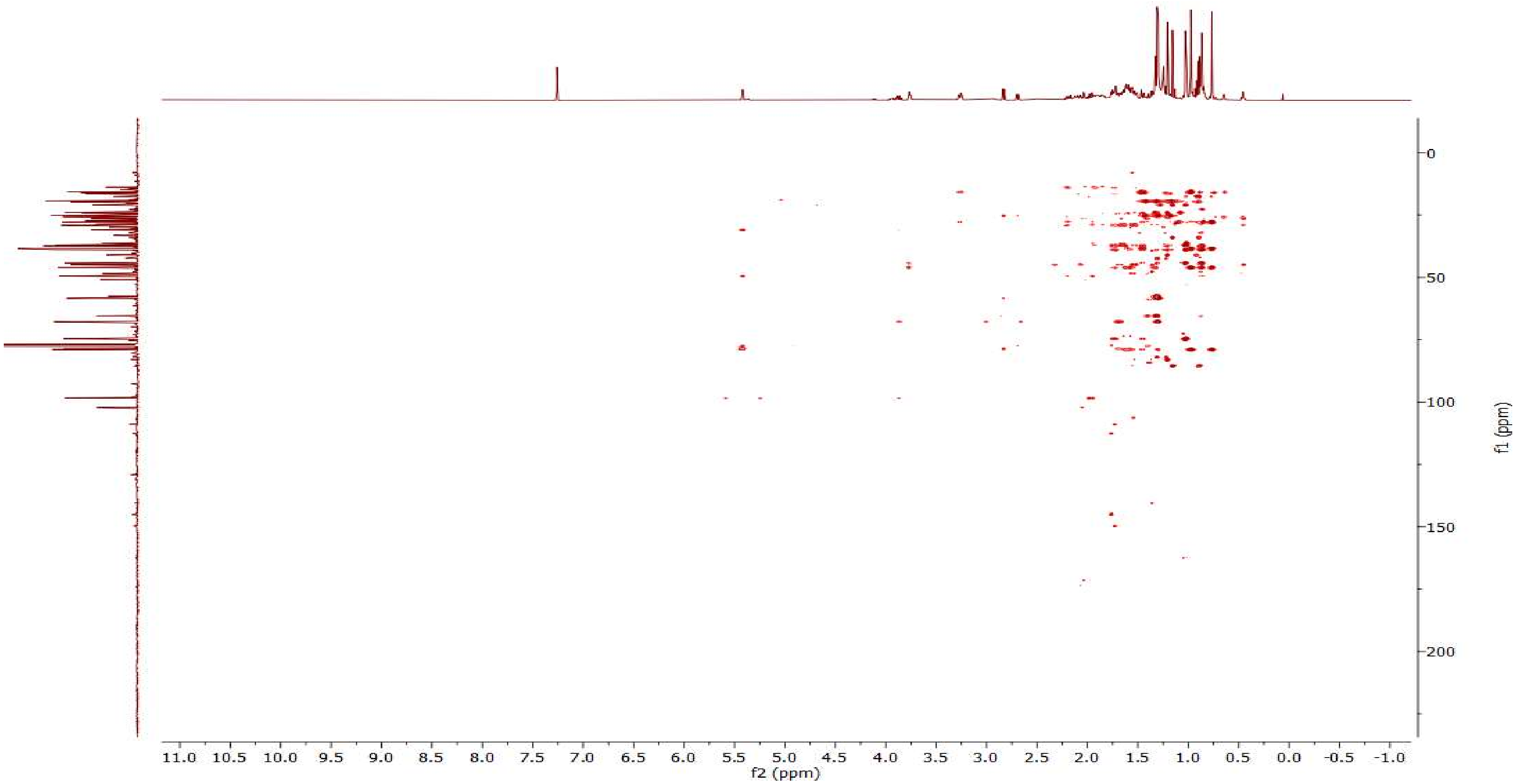


**Figure S21. HMBC spectrum of protoglabretal (11) ($CDCl_3$, 298 K, 500 MHz).**

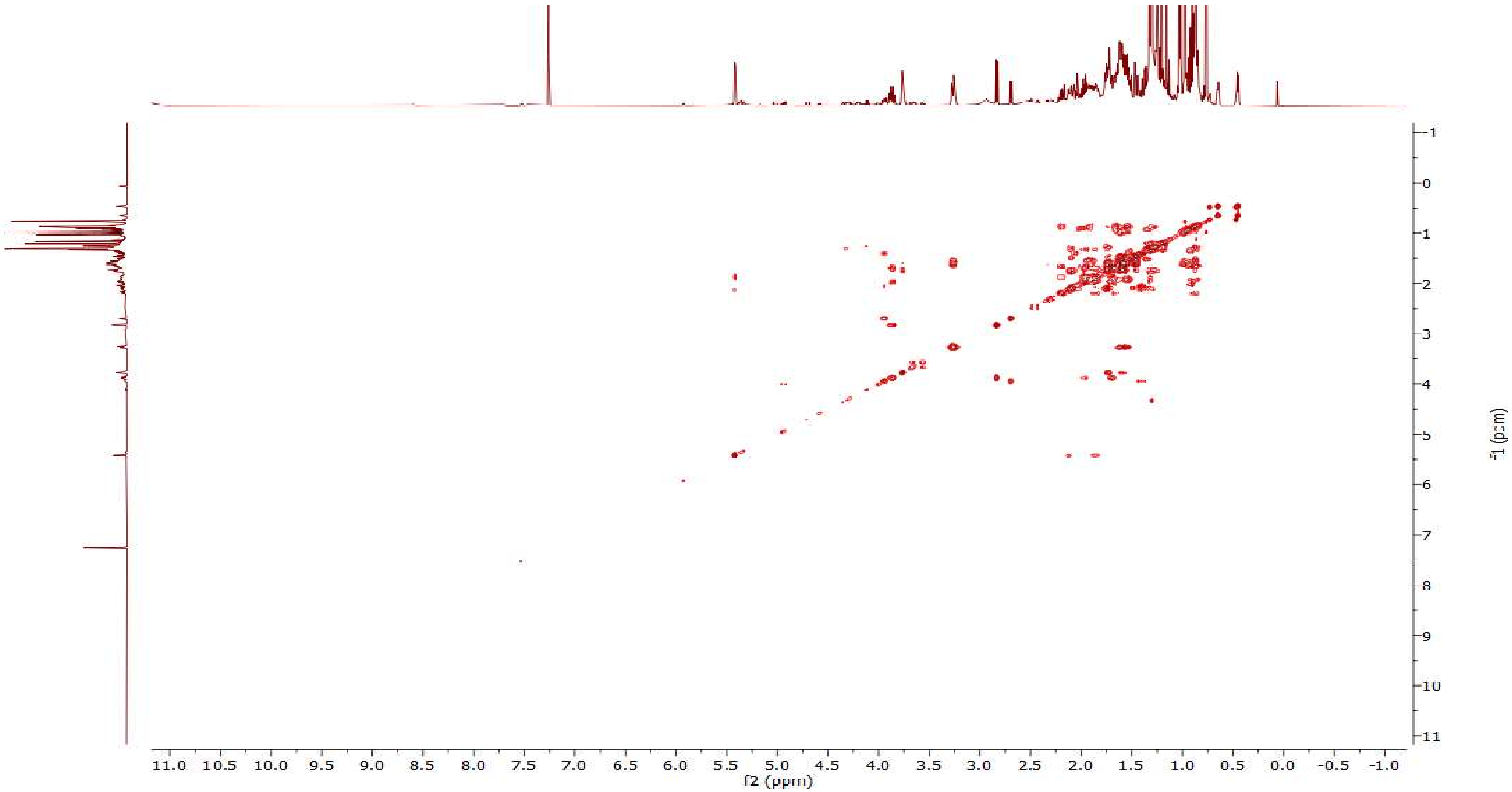


**Figure S22. COSY spectrum of protoglabretal (11) ($CDCl_3$, 298 K, 500 MHz).**

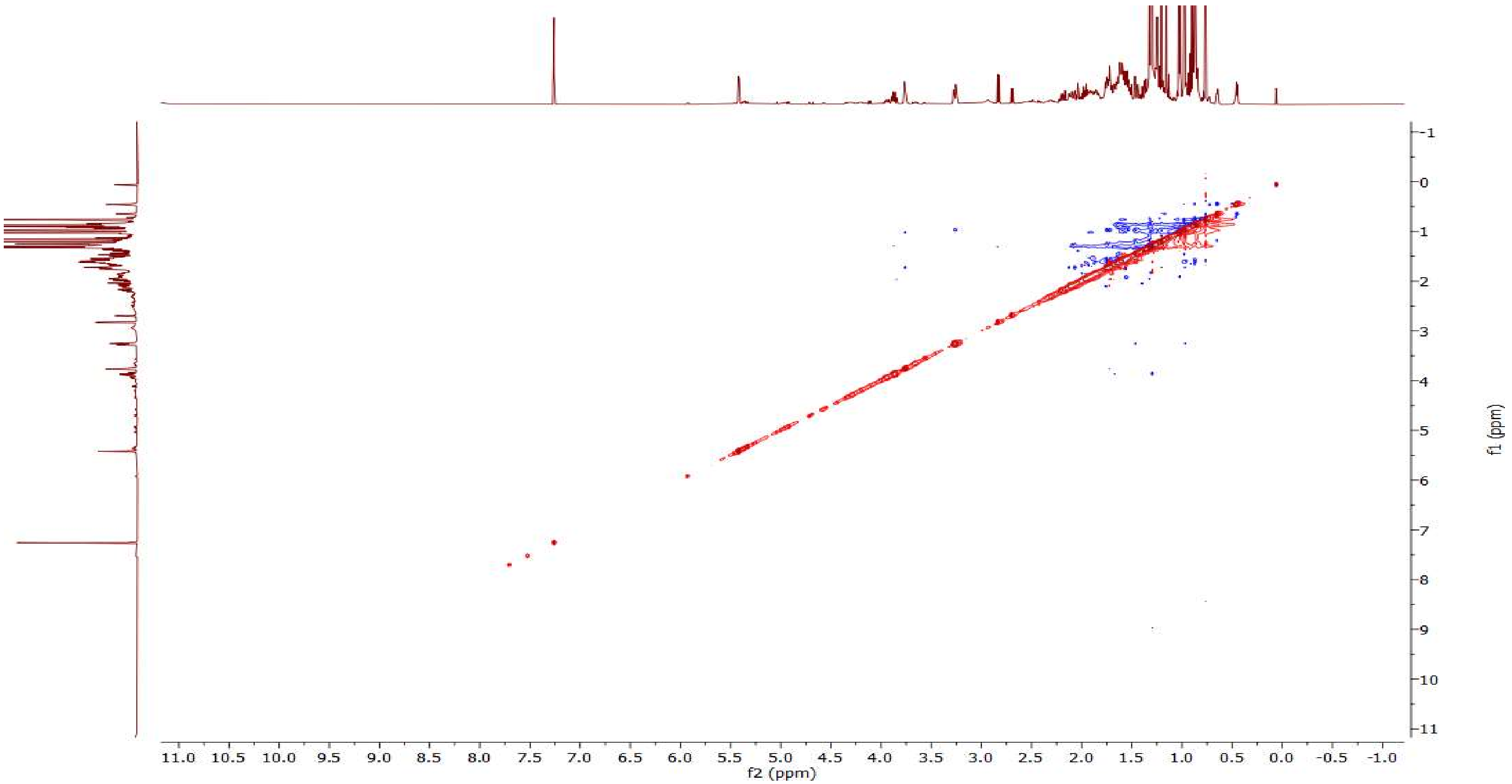


**Figure S23. NOESY spectrum of protoglabretal (11) ($CDCl_3$, 298 K, 500 MHz).**

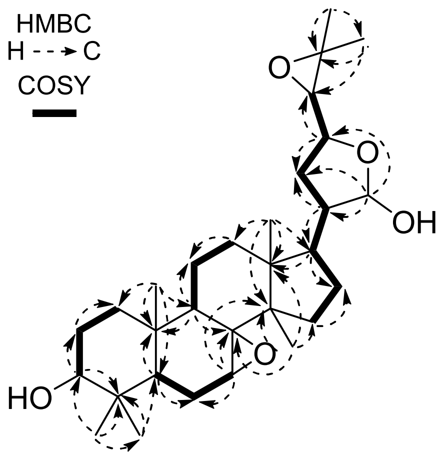


**Figure S24. Selected key HMBC and COSY correlations of 7,8-epoxymelianol (9).**

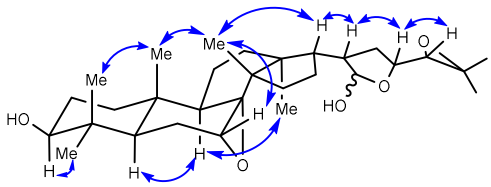


**Figure S25. Selected NOE correlations of 7,8-epoxymelianol (9).**

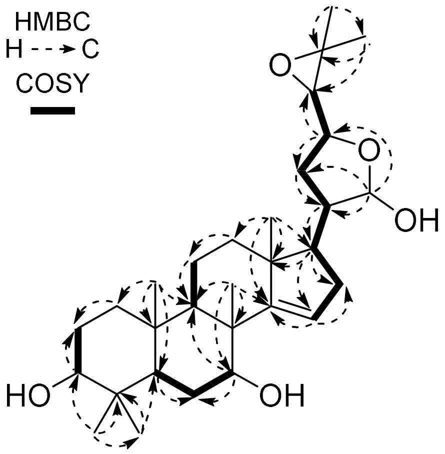


**Figure S26. Selected key HMBC and COSY correlations of isomeliandiol (10).**

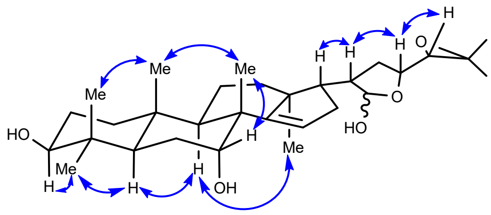


Figure S27. Selected NOE correlations of isomeliandiol (10).

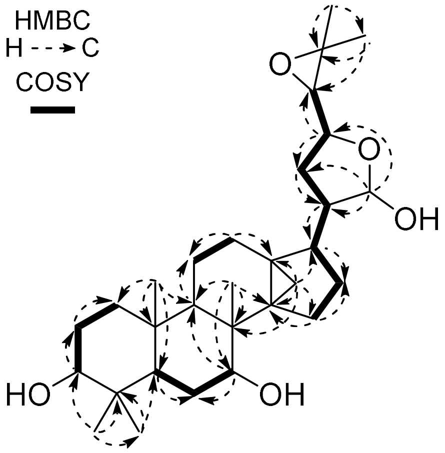


Figure S28. Selected key HMBC and COSY correlations of protoglabretal (11).

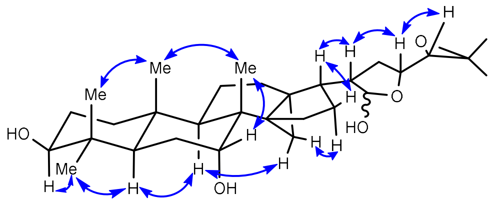


Figure S29. Selected NOE correlations of protoglabretal (11).

**Table S7. Representative examples of isoprotolimonoids previously isolated from Meliaceae, Rutaceae, and Simaroubaceae plants out of ca. 120 structurally related natural products listed in Reaxys.** Stereochemistry is shown as reported in the literature.

| Trivial name | Structure | Family | Subfamily | Species | Reference |
|---|---|---|---|---|---|
| No trivial name | 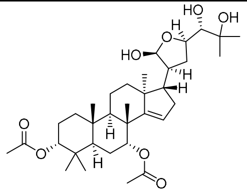 | Meliaceae | Aglaieae | *Aglaia odorata* var. *microphyllina* | [33] |
| Agladoral C | 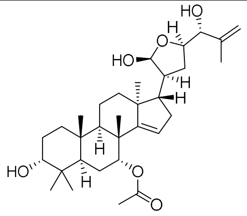 | Meliaceae | Aglaieae | *Aglaia odorata* var. *microphyllina* | [33] |
| Toonasinensin A | 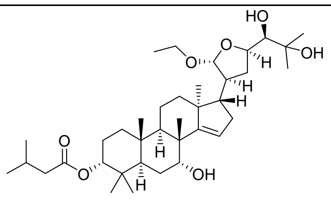 | Meliaceae | Cedreleae | *Toona sinensis* | [34] |
| No trivial name | 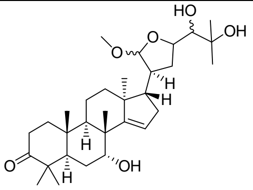 | Meliaceae | Cedreleae | *Cedrela sinensis* | [35] |
| Toonaciliatine A | 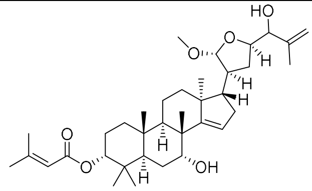 | Meliaceae | Cedreleae | *Toona ciliata* | [36] |
| 21-*O*-Acetyl-toosendantriol | 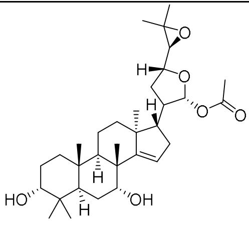 | Meliaceae | Melioideae | *Melia toosendan* | [37] |
| Cumingianol D | 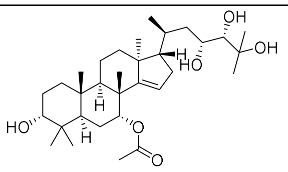 | Meliaceae | Melioideae | *Dysoxylum cumingianum* | [38] |

| | | | | | |
|---|---|---|---|---|---|
| Lepidotrichilin B | 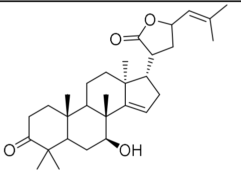 | Meliaceae | Trichilieae | *Trichilia lepidota* | [39] |
| Feroniellide C | 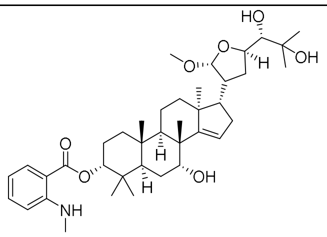 | Rutaceae | Aurantioideae | *Feroniella lucida* | [40] |
| Chisocheton A | 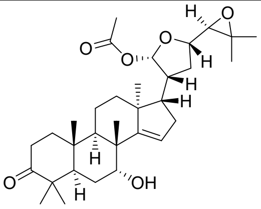 | Rutaceae | Zanthoxyloideae | *Vepris uguenensis* | [41] |
| Brujavanone L | 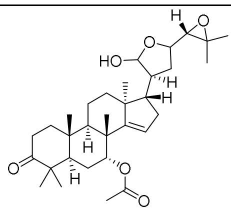 | Simaroubaceae | | *Brucea javanica* | [42] |
| No trivial name | 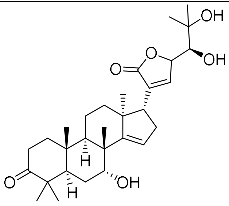 | Simaroubaceae | | *Picrolemma granatensis* | [43] |

**Table S8. Representative examples of glabretanes previously isolated from Meliaceae, Rutaceae, and Simaroubaceae plants out of ca. 110 structurally related natural products listed in Reaxys.** Stereochemistry is shown as reported in the literature.

| Trivial name | Structure | Family | Subfamily | Species | Reference |
|---|---|---|---|---|---|
| 7-Deacetylglabretal-3-acetate | 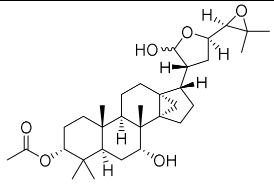 | Meliaceae | Aglaieae | *Aglaia ferruginaea* | [44] |
| No trivial name | 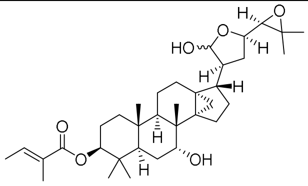 | Meliaceae | Aglaieae | *Aglaia crassinervia* | [45] |
| No trivial name | 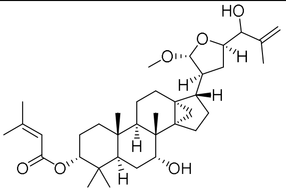 | Meliaceae | Cedreleae | *Cedrela sinensis* | [46] |
| Glabretal | 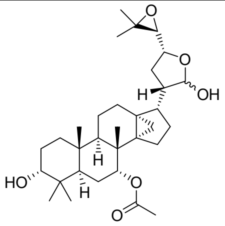 | Meliaceae | Melioideae | *Guarea glabra* | [47] |
| Cumingianoside F | 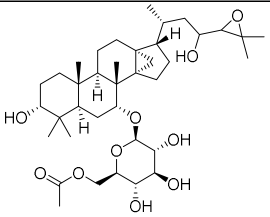 | Meliaceae | Melioideae | *Dysoxylum cumingianum* | [48] |
| Cumingianoside D | 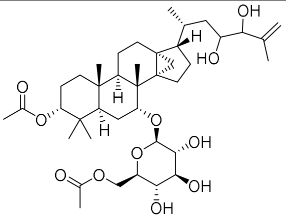 | Meliaceae | Melioideae | *Dysoxylum cumingianum* | [48] |

| | | | | | |
|---|---|---|---|---|---|
| No trivial name | 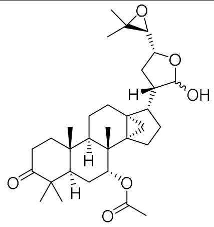 | Meliaceae | Melioideae | *Guarea glabra* | [47] |
| Dysoxylic acid B | 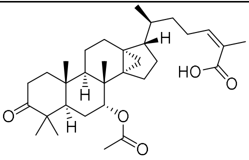 | Meliaceae | Melioideae | *Dysoxylum pettigrewianum* | [49] |
| Dysoxin 2B | 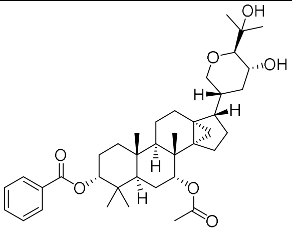 | Meliaceae | Melioideae | *Dysoxylum muelleri* | [50] |
| 3-Episkimmiarepin A | 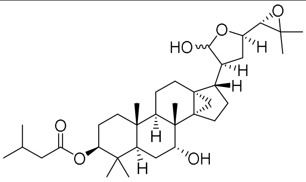 | Rutaceae | Aurantioideae | *Luvunga sarmentosa* | [51] |
| Skimmiarepin A | 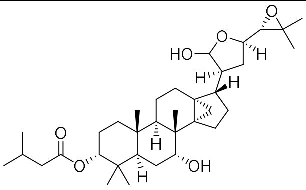 | Rutaceae | Aurantioideae | *Aegle marmelos* | [52] |
| | | Rutaceae | Aurantioideae | *Skimmia japonica* | [53] |
| No trivial name | 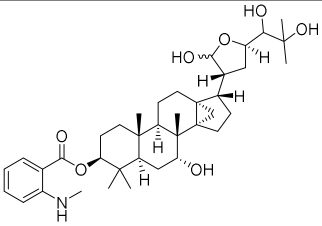 | Rutaceae | Zanthoxyloideae | *Raulinoa echinata* | [54] |
| 3-Oxoskimmiarepin | 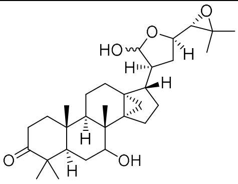 | Rutaceae | Zanthoxyloideae | *Zanthoxylum petiolare* | [55] |

| | | | | | |
|---|---|---|---|---|---|
| Ailanthol | 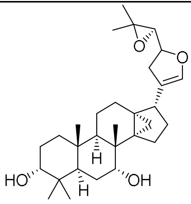 | Simaroubaceae | | *Ailanthus malabarica* | [56,57] |
| Ailanthusin C | 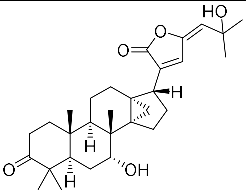 | Simaroubaceae | | *Ailanthus triphysa* | [58] |

**Ergosterol biosynthesis (fungi):**

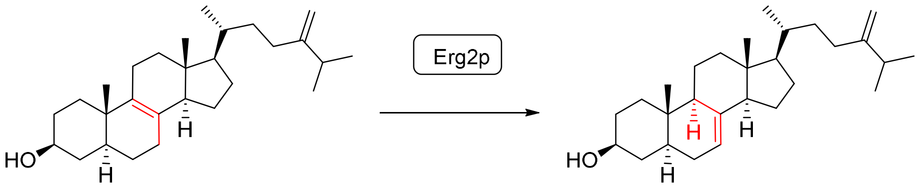


Fecosterol Episterol

**Cholesterol biosynthesis (animals):**

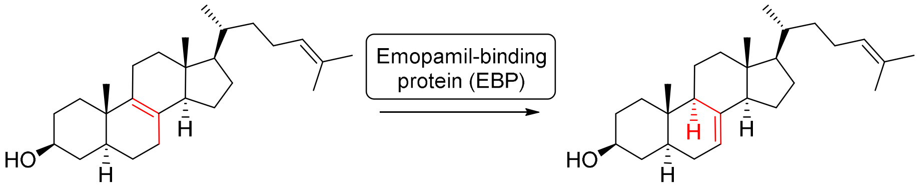


Zymosterol Dehydrolathosterol

**Phytosterol biosynthesis (plants):**

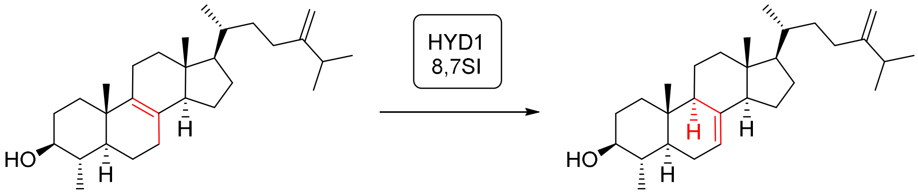


4α-Methylfecosterol 24-Methylenelophenol

**Figure S30. Overview over common C-8,7 sterol isomerase (8,7SI) reactions in primary metabolism.**

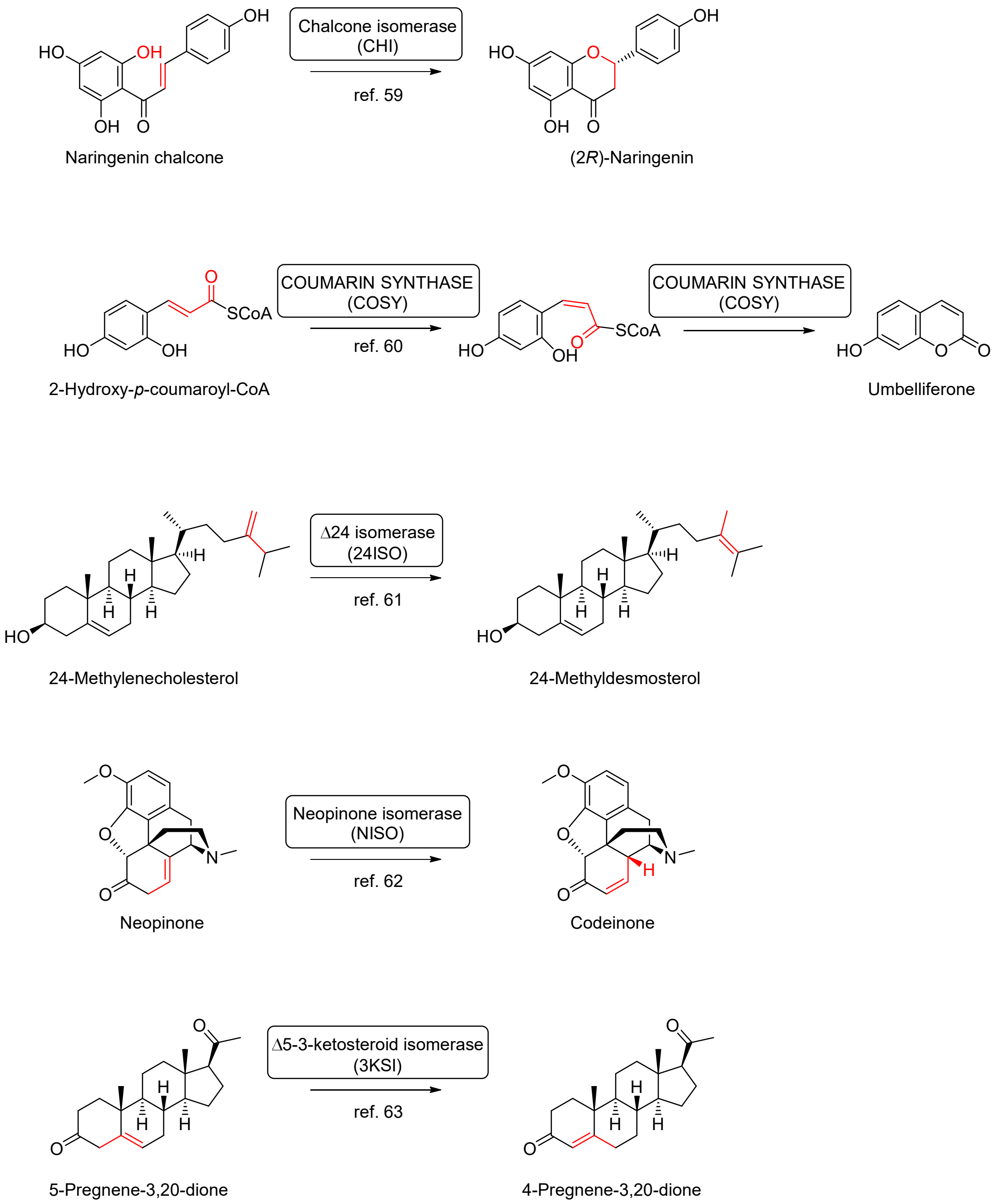


**Figure S31. Overview over other isomerase-catalysed reactions in plant specialised metabolism.**[59–63]

#### References


[1] L. Chuang, S. Liu, D. Biedermann, J. Franke, *Front. Plant Sci.* **2022**, *13*.

[2] J. Bally, H. Jung, C. Mortimer, F. Naim, J. G. Philips, R. Hellens, A. Bombarely, M. M. Goodin, P. M. Waterhouse, *Annu. Rev. Phytopathol.* **2018**, *56*, 405–426.

[3] L. Chuang, J. Franke, in *Eng. Nat. Prod. Biosynth. Methods Protoc.* (Ed.: E. Skellam), Springer US, New York, NY, **2022**, pp. 395–420.

[4] B. J. Haas, A. Papanicolaou, M. Yassour, M. Grabherr, P. D. Blood, J. Bowden, M. B. Couger, D. Eccles, B. Li, M. Lieber, M. D. MacManes, M. Ott, J. Orvis, N. Pochet, F. Strozzi, N. Weeks, R. Westerman, T. William, C. N. Dewey, R. Henschel, R. D. LeDuc, N. Friedman, A. Regev, *Nat. Protoc.* **2013**, *8*, 1494–1512.

[5] R. Patro, G. Duggal, M. I. Love, R. A. Irizarry, C. Kingsford, *Nat. Methods* **2017**, *14*, 417–419.

[6] M. D. Robinson, A. Oshlack, *Genome Biol.* **2010**, *11*, R25.

[7] R. Wehrens, L. M. C. Buydens, *J. Stat. Softw.* **2007**, *21*, 1–19.

[8] R. M. E. Payne, D. Xu, E. Foureau, M. I. S. Teto Carqueijeiro, A. Oudin, T. D. de Bernonville, V. Novak, M. Burow, C.-E. Olsen, D. M. Jones, E. C. Tatsis, A. Pendle, B. A. Halkier, F. Geu-Flores, V. Courdavault, H. H. Nour-Eldin, S. E. O'Connor, *Nat. Plants* **2017**, *3*, 16208.

[9] "Home · TransDecoder/TransDecoder Wiki," can be found under https://github.com/TransDecoder/TransDecoder, **n.d.**

[10] J. Mistry, R. D. Finn, S. R. Eddy, A. Bateman, M. Punta, *Nucleic Acids Res.* **2013**, *41*, e121.

[11] J. Mistry, S. Chuguransky, L. Williams, M. Qureshi, G. A. Salazar, E. L. L. Sonnhammer, S. C. E. Tosatto, L. Paladin, S. Raj, L. J. Richardson, R. D. Finn, A. Bateman, *Nucleic Acids Res.* **2021**, *49*, D412–D419.

[12] F. Sainsbury, E. C. Thuenemann, G. P. Lomonossoff, *Plant Biotechnol. J.* **2009**, *7*, 682–693.

[13] H. Peyret, J. K. M. Brown, G. P. Lomonossoff, *Plant Methods* **2019**, *15*, 108.

[14] J. Reed, M. J. Stephenson, K. Miettinen, B. Brouwer, A. Leveau, P. Brett, R. J. M. Goss, A. Goossens, M. A. O'Connell, A. Osbourn, *Metab. Eng.* **2017**, *42*, 185–193.

[15] A. Horn, U. Kazmaier, *Eur. J. Org. Chem.* **2018**, *2018*, 2531–2536.

[16] R. J. Grebenok, T. E. Ohnmeiss, A. Yamamoto, E. D. Huntley, D. W. Galbraith, D. Della Penna, *Plant Mol. Biol.* **1998**, *38*, 807–815.

[17] Q. Xu, L.-L. Chen, X. Ruan, D. Chen, A. Zhu, C. Chen, D. Bertrand, W.-B. Jiao, B.-H. Hao, M. P. Lyon, J. Chen, S. Gao, F. Xing, H. Lan, J.-W. Chang, X. Ge, Y. Lei, Q. Hu, Y. Miao, L. Wang, S. Xiao, M. K. Biswas, W. Zeng, F. Guo, H. Cao, X. Yang, X.-W. Xu, Y.-J. Cheng, J. Xu, J.-H. Liu, O. J. Luo, Z. Tang, W.-W. Guo, H. Kuang, H.-Y. Zhang, M. L. Roose, N. Nagarajan, X.-X. Deng, Y. Ruan, *Nat. Genet.* **2013**, *45*, 59–66.

[18] X. Wang, Y. Xu, S. Zhang, L. Cao, Y. Huang, J. Cheng, G. Wu, S. Tian, C. Chen, Y. Liu, H. Yu, X. Yang, H. Lan, N. Wang, L. Wang, J. Xu, X. Jiang, Z. Xie, M. Tan, R. M. Larkin, L.-L. Chen, B.-G. Ma, Y. Ruan, X. Deng, Q. Xu, *Nat. Genet.* **2017**, *49*, 765–772.

[19] Y.-T. Ji, Z. Xiu, C.-H. Chen, Y. Wang, J.-X. Yang, J.-J. Sui, S.-J. Jiang, P. Wang, S.-Y. Yue, Q.-Q. Zhang, J. Jin, G.-S. Wang, Q.-Q. Wei, B. Wei, J. Wang, H.-L. Zhang, Q.-Y. Zhang, J. Liu, C.-J. Liu, J.-B. Jian, C.-Q. Qu, *Mol. Ecol. Resour.* **2021**, *21*, 1243–1255.

[20] P. Wang, Y. Luo, J. Huang, S. Gao, G. Zhu, Z. Dang, J. Gai, M. Yang, M. Zhu, H. Zhang, X. Ye, A. Gao, X. Tan, S. Wang, S. Wu, E. B. Cahoon, B. Bai, Z. Zhao, Q. Li, J. Wei, H. Chen, R. Luo, D. Gong, K. Tang, B. Zhang, Z. Ni, G. Huang, S. Hu, Y. Chen, *Genome Biol.* **2020**, *21*, 60.

[21] J. Yang, H. M. Wariss, L. Tao, R. Zhang, Q. Yun, P. Hollingsworth, Z. Dao, G. Luo, H. Guo, Y. Ma, W. Sun, *GigaScience* **2019**, *8*, giz085.

[22] A. Rahier, S. Pierre, G. Riveill, F. Karst, *Biochem. J.* **2008**, *414*, 247–259.

[23] S. Kikuchi, K. Satoh, T. Nagata, N. Kawagashira, K. Doi, N. Kishimoto, J. Yazaki, M. Ishikawa, H. Yamada, H. Ooka, I. Hotta, K. Kojima, T. Namiki, E. Ohneda, W. Yahagi, K. Suzuki, C. J. Li, K. Ohtsuki, T. Shishiki, Y. Otomo, K. Murakami, Y. Iida, S. Sugano, T. Fujimura, Y. Suzuki, Y. Tsunoda, T. Kurosaki, T. Kodama, H. Masuda, M. Kobayashi, Q. Xie, M. Lu, R. Narikawa, A. Sugiyama, K. Mizuno, S. Yokomizo, J. Niikura, R. Ikeda, J. Ishibiki, M. Kawamata, A. Yoshimura, J. Miura, T. Kusumegi, M. Oka, R. Ryu, M. Ueda, K. Matsubara, J. Kawai, P. Carninci, J. Adachi, K. Aizawa, T. Arakawa, S. Fukuda, A. Hara, W. Hashidume, N. Hayatsu, K. Imotani, Y. Ishii, M. Itoh, I. Kagawa, S. Kondo, H. Konno, A. Miyazaki, N. Osato, Y. Ota, R. Saito, D. Sasaki, K. Sato, K. Shibata, A. Shinagawa, T. Shiraki, M. Yoshino, Y. Hayashizaki, *Science* **2003**, *301*, 376–379.

[24] S.-H. Bae, J. N. Lee, B. U. Fitzky, J. Seong, Y.-K. Paik, *J. Biol. Chem.* **1999**, *274*, 14624–14631.

[25] S. Silve, P. H. Dupuy, C. Labit-Lebouteiller, M. Kaghad, P. Chalon, A. Rahier, M. Taton, J. Lupker, D. Shire, G. Loison, *J. Biol. Chem.* **1996**, *271*, 22434–22440.

[26] M. Hanner, F. F. Moebius, F. Weber, M. Grabner, J. Striessnig, H. Glossmann, *J. Biol. Chem.* **1995**, *270*, 7551–7557.

[27] T. Long, A. Hassan, B. M. Thompson, J. G. McDonald, J. Wang, X. Li, *Nat. Commun.* **2019**, *10*, 2452.

[28] H. Yao, H. Cai, D. Li, *J. Mol. Biol.* **2020**, *432*, 5162–5183.

[29] R. C. Edgar, S. Batzoglou, *Curr. Opin. Struct. Biol.* **2006**, *16*, 368–373.

[30] F. Sievers, D. G. Higgins, *Protein Sci.* **2018**, *27*, 135–145.

[31] F. Sievers, A. Wilm, D. Dineen, T. J. Gibson, K. Karplus, W. Li, R. Lopez, H. McWilliam, M. Remmert, J. Söding, J. D. Thompson, D. G. Higgins, *Mol. Syst. Biol.* **2011**, *7*, 539.

[32] S. Guindon, J.-F. Dufayard, V. Lefort, M. Anisimova, W. Hordijk, O. Gascuel, *Syst. Biol.* **2010**, *59*, 307–321.

[33] J. Liu, S.-P. Yang, G. Ni, Y.-C. Gu, J.-M. Yue, *J. Asian Nat. Prod. Res.* **2012**, *14*, 929–939.

[34] D. Liu, R. Wang, L. Xuan, X. Wang, W. Li, *Molecules* **2020**, *25*, 801.

[35] K. Mitsui, H. Saito, R. Yamamura, H. Fukaya, Y. Hitotsuyanagi, K. Takeya, *Chem. Pharm. Bull. (Tokyo)* **2007**, *55*, 1442–1447.

[36] J. Ning, H.-P. He, S.-F. Li, Z.-L. Geng, X. Fang, Y.-T. Di, S.-L. Li, X.-J. Hao, *J. Asian Nat. Prod. Res.* **2010**, *12*, 448–452.

[37] T. Nakanishi, A. Inada, M. Nishi, T. Miki, R. Hino, T. Fujiwara, *Chem. Lett.* **1986**, *15*, 69–72.

[38] S. Kurimoto, Y. Kashiwada, K.-H. Lee, Y. Takaishi, *Phytochemistry* **2011**, *72*, 2205–2211.

[39] W. D. S. Terra, I. J. C. Vieira, R. Braz-Filho, W. R. de Freitas, M. M. Kanashiro, M. C. M. Torres, *Molecules* **2013**, *18*, 12180–12191.

[40] P. Phuwapraisirisan, S. Sombund, S. Tip-pyang, P. Siripong, *Nat. Prod. Res.* **2013**, *27*, 753–760.

[41] J. J. Kiplimo, Md. Shahidul Islam, N. A. Koorbanally, *Phytochemistry* **2012**, *83*, 136–143.

[42] S.-H. Dong, J. Liu, Y.-Z. Ge, L. Dong, C.-H. Xu, J. Ding, J.-M. Yue, *Phytochemistry* **2013**, *85*, 175–184.

[43] E. R. Fo, J. B. Fernandes, P. C. Vieira, M. F. das G. F. da Silva, J. Zukerman-Schpector, R. M. O. Corrêa de Lima, S. C. Nascimento, W. Thomas, *Phytochemistry* **1996**, *43*, 857–862.
[44] D. A. Mulholland, T. V. Monkhe, *Phytochemistry* **1993**, *34*, 579–580.
[45] B.-N. Su, H. Chai, Q. Mi, S. Riswan, L. B. S. Kardono, J. J. Afriastini, B. D. Santarsiero, A. D. Mesecar, N. R. Farnsworth, G. A. Cordell, S. M. Swanson, A. D. Kinghorn, *Bioorg. Med. Chem.* **2006**, *14*, 960–972.
[46] K. Mitsui, M. Maejima, H. Saito, H. Fukaya, Y. Hitotsuyanagi, K. Takeya, *Tetrahedron* **2005**, *61*, 10569–10582.
[47] G. Ferguson, P. A. Gunn, W. C. Marsh, R. McCrindle, R. Restivo, J. D. Connolly, J. W. B. Fulke, M. S. Henderson, *J. Chem. Soc. Chem. Commun.* **1973**, 159–160.
[48] Y. Kashiwada, T. Fujioka, J. J. Chang, I. S. Chen, K. Mihashi, K. H. Lee, *J. Org. Chem.* **1992**, *57*, 6946–6953.
[49] D. A. Mulholland, J. J. Nair, *Phytochemistry* **1994**, *37*, 1409–1411.
[50] D. A. Mulholland, J. J. Nair, D. A. H. Taylor, *Phytochemistry* **1996**, *42*, 1667–1671.
[51] C. Kamperdick, T. P. Lien, G. Adam, T. V. Sung, *J. Nat. Prod.* **2003**, *66*, 675–678.
[52] J. Li, F. Mahdi, L. Du, S. Datta, D. G. Nagle, Y.-D. Zhou, *J. Nat. Prod.* **2011**, *74*, 1894–1901.
[53] M. Ochi, A. Tatsukawa, N. Seki, H. Kotsuki, K. Shibata, *Bull. Chem. Soc. Jpn.* **1988**, *61*, 3225–3229.
[54] M. W. Biavatti, P. C. Vieira, M. F. G. F. da Silva, J. B. Fernandes, S. Albuquerque, *J. Nat. Prod.* **2002**, *65*, 562–565.
[55] M. S. P. Arruda, J. B. Fernandes, P. C. Vieira, M. Fatima Das G.F. Da Silva, J. R. Pirani, *Phytochemistry* **1994**, *36*, 1303–1306.
[56] B. S. Joshi, V. N. Kamat, S. William Pelletier, K. Go, K. Bhandary, *Tetrahedron Lett.* **1985**, *26*, 1273–1276.
[57] Y. Hitotsuyanagi, A. Ozeki, C. Y. Choo, K. L. Chan, H. Itokawa, K. Takeya, *Tetrahedron* **2001**, *57*, 7477–7480.
[58] S. Thongnest, J. Boonsombat, H. Prawat, C. Mahidol, S. Ruchirawat, *Phytochemistry* **2017**, *134*, 98–105.
[59] J. M. Jez, M. E. Bowman, R. A. Dixon, J. P. Noel, *Nat. Struct. Biol.* **2000**, *7*, 786–791.
[60] R. Vanholme, L. Sundin, K. C. Seetso, H. Kim, X. Liu, J. Li, B. D. Meester, L. Hoengenaert, G. Goeminne, K. Morreel, J. Haustraete, H.-H. Tsai, W. Schmidt, B. Vanholme, J. Ralph, W. Boerjan, *Nat. Plants* **2019**, *5*, 1066–1075.
[61] E. Knoch, S. Sugawara, T. Mori, C. Poulsen, A. Fukushima, J. Harholt, Y. Fujimoto, N. Umemoto, K. Saito, *Proc. Natl. Acad. Sci. U. S. A.* **2018**, *115*, E8096–E8103.
[62] M. Dastmalchi, X. Chen, J. M. Hagel, L. Chang, R. Chen, S. Ramasamy, S. Yeaman, P. J. Facchini, *Nat. Chem. Biol.* **2019**, *15*, 384–390.
[63] N. Meitinger, D. Geiger, T. W. Augusto, R. Maia de Pádua, W. Kreis, *Phytochemistry* **2015**, *109*, 6–13.
